# Ant head shape evolved to compromise bite-induced mechanical demands

**DOI:** 10.1101/2025.03.19.643247

**Authors:** Cristian L. Klunk, Alexandre Casadei-Ferreira, Marco A. Argenta, Marcio R. Pie

## Abstract

Heads of ant workers harbour the mouthparts and respective musculature, varying drastically in morphology. The mandible adductor muscles occupy most of the head’s internal volume, and their contraction generates forces that could risk cuticle failure. Here we quantified ant worker plane head shape disparity and explored how it influences stress dissipation under biting conditions. We combined a geometric morphometric approach under a phylogenetic comparative framework and biomechanical simulations to test the hypothesis that idealized head shapes poorly explored by current ant lineages exhibit biomechanical limitations that could be negatively selected, while an idealized shape representing the median ant worker head shape would present a superior mechanical performance. Our results revealed that narrow heads with deep vertex depressions distribute stresses more evenly, but usually at high levels. Broader heads with convex posterior margins, in contrast, concentrate stresses around the mandibular articulations, making them more prone to failure under high bite forces. Most ant lineages exhibit a head shape that allows mild stresses to spread along the head while concentrating higher stresses around the mandibular articulations. This evolutionary pattern results in a limited morphospace occupation, with most ants clustering around a typical shape and only a few lineages exploring extreme morphologies.

## Introduction

Morphological evolution is commonly associated with functional aspects (Koehl 1996), and there are instances where functional adaptation demands a degree of morphological specialization that limits its variability (McGhee 2015), usually due to trade-offs in performance that hamper exploration of novel morphologies. In other circumstances, however, morphological diversity can converge functionally, an evolutionary phenomenon known as “many-to-one mapping” (Wainwright 2005, 2007). A classic example of a close association between a morphological structure and its function is the beak of Darwin finches, whose diet varies in the size and hardness of food items matched by adaptations on beak shape (Soons *et al*. 2015). Functional morphology, therefore, represents a widely explored research avenue to investigate potential evolutionary pressures that explain the extent and limitations of morphological evolution.

The remarkable diversity in ant worker head and mandible morphology suggests that functional aspects could have influenced their morphological evolution. Ant workers rely on their mandibles to execute most colony tasks, involving behaviours like biting, excavating, carrying, cutting, and fighting (Wheeler 1910, Wilson 1987). Indeed, the versatility of ant mandibles potentially represents one of the main characteristics of their ecological success (Wilson 1987). Two muscle pairs located inside the head capsule perform the mandibular movement. However, the mandibular adductor muscles (0md1), responsible for mandible closing, are much more developed and occupy most of the head’s internal volume (Paul 2001, Paul and Gronenberg 2002, Lillico-Ouachour *et al*. 2018, Richter *et al*. 2019, 2020, 2021, Boudinot *et al*. 2021, Püffel *et al*. 2021).

Head shape affects muscle packing, and relative differences in growing patterns between head regions represent a relevant component of ant morphological evolution (Pie and Traniello 2007, Pie and Tschá 2013). It has been suggested that disproportional increases in head width could allow for the accommodation of larger 0md1 muscles (Püffel *et al*. 2021, 2023a, Paul and Gronenberg 1999, Paul 2001) without the necessity to increase the head along its remaining axes (i.e. length and height). Proportional increases in all head dimensions could lead to the development of a significantly heavier head, which could impair body equilibrium (Anderson *et al*. 2020) or represent outstanding amounts of resources for its development (Peeters and Ito 2015, Püffel *et al*. 2021, 2023b). The ant head capsule also provides space for other muscle groups (e.g., antennal and sucking pump muscles), the remaining feeding apparatus (e.g., mouth parts appendages and sucking pump), as well as several glands, along with the central nervous system (Richter *et al*. 2019, 2020, 2021, Richter and Economo 2023). Studies on ant head evolution provide detailed descriptions of the head’s internal and external anatomy (e.g., Boudinot *et al*. 2021, Richter *et al*. 2022), as well as contributing to our understanding of the degree of modularity and integration in the evolution of head morphology regarding other body regions, like the mandibles (Casadei-Ferreira *et al*. 2021) and mesosoma (Friedman *et al*. 2020), which can represent possible development constraints that limit the range of head shape variation in ants.

In addition to space for muscle packing, the ant worker head is also relevant in other functional contexts. Specialized workers in the ant genus *Cephalotes* use their heads to plug their nest entrance, and the collective strategy adopted to seal the nest depends on the head shape of those workers (Powel *et al*. 2020). Mandibles articulate with the head capsule, whose morphology influences the establishment of bite power amplification mechanisms in *Strumigenys* ants (Booher *et al*. 2021). The head capsule also offers mechanical support for muscle contraction, which result in relevant mechanical demands on the head cuticle (Blanke *et al*. 2017, 2018). Along with the direct mechanical effects of mandibular closing muscle contraction in the head capsule, biting also adds another mechanical impact through the reaction forces generated at the mandibular articulations with the head (Blanke *et al*. 2017). The combined effects of muscle contraction and reaction forces result in the generation of stresses on the head capsule, whose dissipation patterns can be influenced by the shape of the head, as suggested for *Pheidole* ants (Klunk *et al*. 2023).

Here, we tested the biomechanical performance of ant worker heads while quantifying their morphological variation. We built a phylomorphospace of ant worker plane-head shape using geometric morphometrics, considering representatives of most extant lineages, and carried out biting simulations on idealized head shapes representing the extents of the phylomorphospace. Although overlooking the full complexity of a structure, the use of 2D data in biomechanical simulations can effectively approximate the mechanical demands of complex biological structures (Marcé-Nogué *et al*. 2013). Such effectiveness was demonstrated by the effects of biting behaviour on skulls of crocodilians (Pierce *et al*. 2008, 2009), theropods (Rayfield 2005) and *Tyrannosaurus rex* (Rayfield 2004), bite loading in vertebrate jaws (Deaking *et al*. 2022), the mechanical consequences of burrowing on the trilobite cephalon (Esteve *et al*. 2021), as well as the effects of biting on *Pheidole* worker heads (Klunk *et al*. 2023).

We hypothesize that the variation in ant head shape will be closely related to head width, length, and the degree of vertex depression, given that these were primary aspects concerning head morphological variation in one of the most diverse ant genera, *Pheidole* (Casadei-Ferreira *et al*. 2021, 2022). We expect that head shapes located in the limits of the phymorphospace will show biomechanical limitations that potentially prevent their evolution or a widely spread among current ant lineages. Accordingly, we also suspect that the most explored head shape represents a structure optimized to deal with the mechanical demands of bite, favouring stress dissipation while avoiding high-level stress concentration on limited regions of the head. These biomechanical hypotheses stand on previous findings regarding *Pheidole* worker heads (Klunk *et al*. 2023).

## Materials and Methods

### Data collection

We downloaded high-resolution images of worker head of several ant species from the online repository antweb.org (AntWeb 2021). Although ant heads are complex, three-dimensional structures with intricate details in the internal and external morphology, the inclusion of all those aspects might lead to substantial methodological and computational challenges, and would drastically reduce sample sizes. Therefore, as a first approximation, we only focused on the two-dimensional contour of the head in full-face view to explore the role of morphological variation in the mechanical responses of these structures to bite-loading. We selected head images of up to three species for each genus whenever those images were adequate to visualize the position of the landmarks and semi-landmarks. When pictures of more than three species were available, we followed the alphabetic order within the genus. In total, we collated images from 765 species (∼5.3% of known species), representing 322 genera (94% of known genera) and all 16 extant subfamilies (Bolton, 2023; Supplementary File 1).

### Geometric Morphometrics (GM)

To describe the head shape, we digitized four landmarks and 50 semi-landmarks (25 on each head side) using tpsDig v.2.31 (Rohlf 2017) (Fig. 1a). Landmarks L1 and L2 represent the left and right limits between head lateral margins and clypeus, L3 represents the midpoint on the clypeus posterior margin, and L4 the midpoint of the vertex margin. Semi-landmarks were used to describe the head left and right sides from L1/L2 to L3 (Fig. 1a). Geometric morphometrics analyses were performed in R version 4.1.0 (R Core Team 2022) using the geomorph package v. 4.0.4 (Baken *et al*. 2021). We used Procrustes superimposition to remove differences in scale, translation, and rotation. The least-squares criterion (Rohlf and Slice 1990) was applied, allowing semi-landmarks to slide between fixed landmarks. Left- and right-side landmarks and semi-landmarks were averaged to remove any effect of bilateral asymmetry. Principal component analysis (PCA) was used to extract the main trends in head shape variation. To account for shared evolutionary history, we generated a phylomorphospace based on the phylogeny published by Divieso *et al*. (2020), encompassing most current ant lineages (Moreau and Bell 2013, Ward *et al*. 2015). Raw coordinate data was pruned using the GEIGER package v. 2.0.10 (Pennell *et al*. 2014), and the mean shape was calculated for each genus using GEOMORPH to match the phylogeny. This pruning procedure resulted in the removing of three subfamilies (Agroecomyrmecinae, Apomyrminae, and Aneuretinae) and 117 genera from the phylomorphoscae in relation to the complete dataset, resulting in a final number of 205 genera considered for the generation of the phylomorphospace. Finally, phylogeny was projected onto the PCA to produce the phylomorphospace (Sidlauskas 2008). To visualize shape variation, we placed distorted shapes into the background of the phylomorphospace following Olsen (2017). These distorted shapes were subsequently used for the mechanical simulations. Therefore, they were named after their positioning along the phylomorphospace axes (PC1 and PC2), from the negative (min) towards the positive (max) edges of each axis, passing through their middle points (median), resulting in an identification that corresponds to the exact location of the head shape on the phylomorphospace (e.g. PC1maxPC2median - positive edge of PC1 and middle position of PC2; Fig.1c).

**Fig.1.**
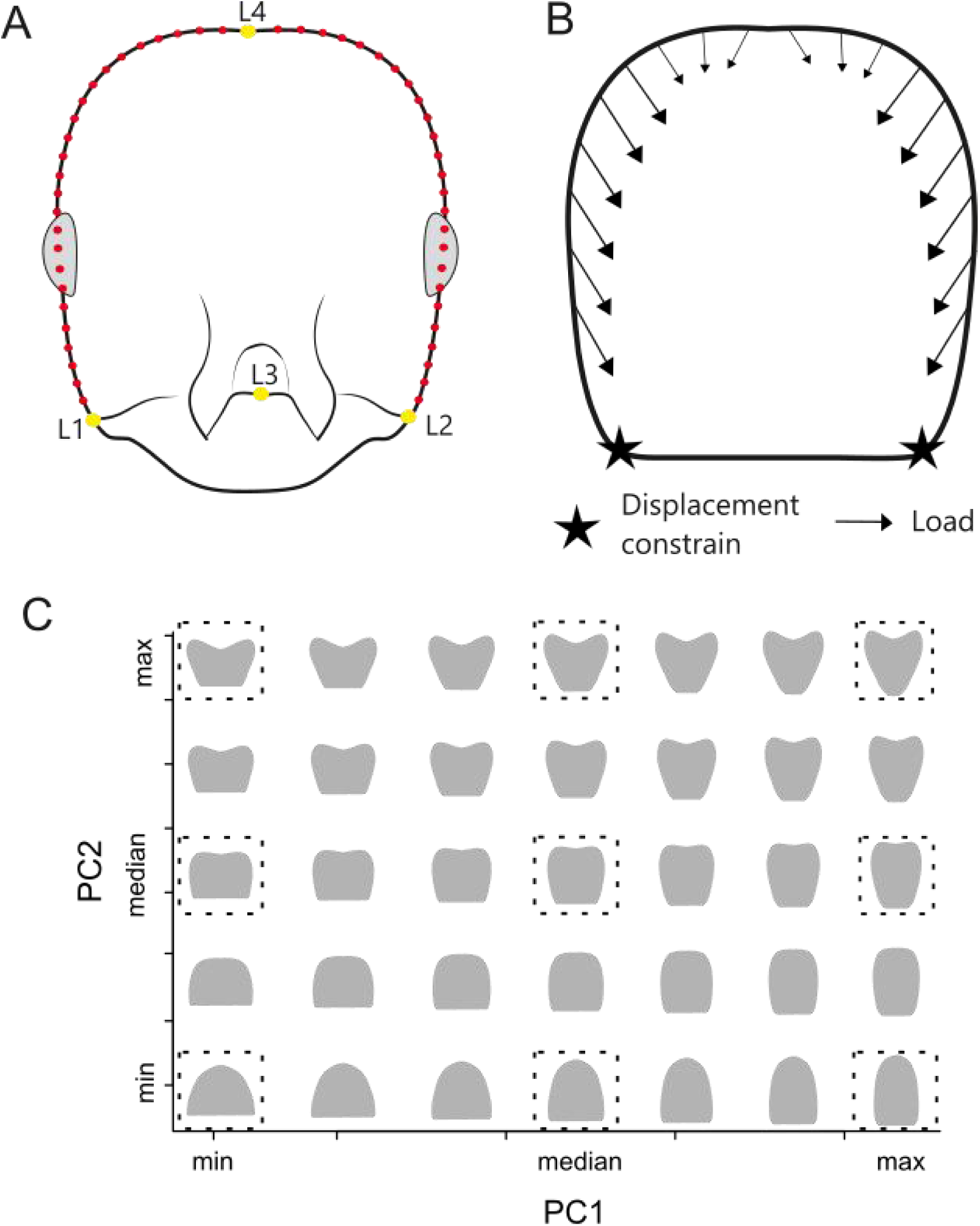
(A) Diagram depicting landmarks (yellow dots) and semi-landmarks (red dots) distribution used to quantify the morphological variation in ant worker heads; (B) placement and direction of loaded and fixed regions (displacement constrains) on the worker head using for FEA; (C) idealized plane head shapes of the morphospace, with dashed squares highlighting the nine head shapes employed in FEA.

### Finite Element Analysis (FEA)

We applied FEA to investigate the influence of plane head shape variation in the mechanical demands from biting, encompassing the effects of 0md1 contraction and the mandibular reaction forces. We considered nine idealized plane head shapes representing distinct morphospace regions (Fig.1c), which were generated using the thin-plate spline deformation grids approach (Bookstein 1991) in R v. 4.1.0 (R Core Team 2022). We vectorized the head pictures in the software Inkscape, meshed them in Fusion 360 (AUTODESK), and then imported them into the finite element solver Abaqus 6 (Dassault Systèmes), where sharp edges were smoothed and plane geometries defined.

We performed linear static simulations with a plane stress approach, which considers that the structure has two main dimensions and the stresses in the third dimension are negligible (Marcé-Nogué *et al*. 2013). Therefore, we applied a unitary and constant thickness to each plane head shape. We designed the finite element meshes with plane triangular and quadratic elements (CPS6M), maintaining a similar mesh density for each model (Table 1). We defined Young’s modulus of the head cuticle as 2.75 GPa, according to measurements taken from leaf-cutting ant mandibles (Brito *et al*. 2017), and the Poisson ratio as 0.3, as commonly considered for ant cuticle (Larabee *et al*. 2018, Zhang *et al*. 2020, Klunk *et al*. 2021, Wang *et al*. 2022). We considered the material properties of the head cuticle as isotropic and linearly elastic.

**Table 1.** Position in the morphospace, mesh area, number of elements and load applied on each head side of the plane head models employed for FEA.

| Morphospace position | Mesh area (mm<sup>2</sup>) | Number of elements | Load (N) |
| --- | --- | --- | --- |
| PC1maxPC2max | 2.16 | 73119 | 0.84 |
| PC1maxPC2mean | 3.03 | 73578 | 1.00 |
| PC1maxPC2min | 2.72 | 72536 | 0.95 |
| PC1meanPC2max | 1.83 | 71785 | 0.78 |
| PC1meanPC2mean | 2.43 | 73708 | 0.90 |
| PC1meanPC2min | 2.43 | 71781 | 0.90 |
| PC1minPC2max | 1.44 | 71990 | 0.70 |
| PC1minPC2mean | 1.85 | 71110 | 0.78 |
| PC1minPC2min | 1.52 | 71986 | 0.71 |

We simulated the contraction of the 0md1 by applying the load at the nodes of the posterior and lateral margins of each head side, with load orientation approximating the main direction of 0md1 muscular fibers (Fig. 1b; Muscedere *et al*. 2011, Lillico-Ouachour *et al*. 2018, Richter *et al*. 2019, 2020, 2021, Püffel *et al*. 2023a). A 1N load was applied in each head side in the model with the largest surface area (PC1maxPC2mean), and the loading magnitude of the remaining models was corrected based on the difference in surface area from that reference model (Table 1; Marcé-Nogué *et al*. 2013). To simulate the bite reaction forces exercised by the mandibles on the head, we fixed at zero the nodal displacement along the x and y directions on head regions that correspond to the regions of mandibular articulation, fixing the same number of nodes for each head representation (Fig. 1b).

We considered two plot types to analyze stress patterns: tensor plots and color maps. Tensor plots depict stress distribution as arrows that indicate the normalized stress value (arrow size), stress direction (arrow orientation), and stress type (compressive with inward-pointed arrowheads, tensile with outward-pointed arrowheads). We considered minimum and maximum principal stresses in tensor plots to show shearless principal normal stress patterns. Color maps depict stress patterns by considering a unique stress value for each mesh element based on a stress transformation criterion. We chose the Tresca criterion to define stress values, which is a criterion employed to predict yielding in ductile materials, being more conservative in its predictions than other criteria usually applied to transform stress values regarding ductile materials, like von Mises and Rankine (Özkaya *et al*. 2017).

### Intervals method

We applied the intervals method to evaluate how worker plane head shapes differ in the amount of area covered by distinct ranges of stress values (Marcé-Nogué *et al*. 2017). This approach defines stress intervals based on the element stress values of each simulated meshes. Since we can extract the area of each plane element, we can quantify the proportion of head area covered by each stress interval. These proportions are then used for further statistical analysis or visualization techniques, such as PCA, to visualize head shape differences in the distribution of non-normalized stresses (Marcé-Nogué *et al*. 2017).

To perform the intervals method, we extracted element Tresca equivalent stress and area values from Abaqus 6 (Dassault Systèmes) and removed elements of the 2% highest stress values for each simulation since they usually represent artificial high values (Marcé-Nogué *et al*. 2016, 2017). We log-transformed stress values before generating the stress intervals to reduce the high variation between extremes of stress values due to the proximity of some elements to the regions of load application and constrained nodes submitted to reaction forces. To designate the stress ranges for each stress interval, we defined the upper threshold as 0.77, so that only 2% of the highest stress values from all simulations were contained in the interval of highest stress (Marcé-Nogué *et al*. 2017). We generated datasets with distinct numbers of intervals (5, 10, 15, 25, 50, and 75) and performed a PCA with each dataset to define the ideal number of stress intervals. The scores of PC1 and PC2 of each dataset were used as variables in linear regressions with the scores of equivalent PCs of the next interval (e.g., PC_1_5intervals ∼ PC_1_15intervals), and the coefficient of determination (R^2^) was considered to analyze the convergence of PC scores (Marcé-Nogué *et al*. 2017). Convergence is achieved when R^2^ cease to increase, and then the dataset with the lower number of intervals is chosen to proceed with the analysis (Marcé-Nogué *et al*. 2017). Convergence occurred with the transition from 15 to 25 intervals, and we will discuss the results from the PCA with 15 stress intervals. To perform the PCA we employed the R packages FactoMineR version 2.4 (Lê *et al*. 2008) and factoextra version 1.0.7.999 (Kassambara and Mundt 2020).

## Results

### Geometric morphometrics

The first two axes of the PCA summarizing ant head morphospace explained 56% and 22% of the variance in the dataset, respectively (Fig.2). The first PC described variation in head length and width, with positive scores representing increasingly narrow and longer heads (Fig.2). The second PC was associated with a change in the curvature of the posterior margin of the head, from a concave posterior margin in its negative range towards a convex posterior margin with a deep vertex depression on its positive range (Fig.2). Most head shapes from current ant genus occupied the lower left region of the morphospace, being represented by a plane head shape broader than longer, showing a nearly straight posterior margin of the head (Fig.2). Most subfamilies (e.g., Pseudomyrmicinae, Myrmiciinae, and Ectatomminae) tended to occupy a restricted region of morphospace. On the other hand, some subfamilies showed a broader morphospace occupation. Myrmicinae, Ponerinae, and Dorylinae covered a wide range of PC1. In contrast, representatives of Myrmicinae and Formicinae covered PC2 more extensively (Fig.2). Myrmicinae is notable due to the placement of several of its genera at the extremes of PC1 and PC2. Consequently, all other ant subfamilies are situated within the distributional range of Myrmicinae (Fig.2).

**Fig.2.**
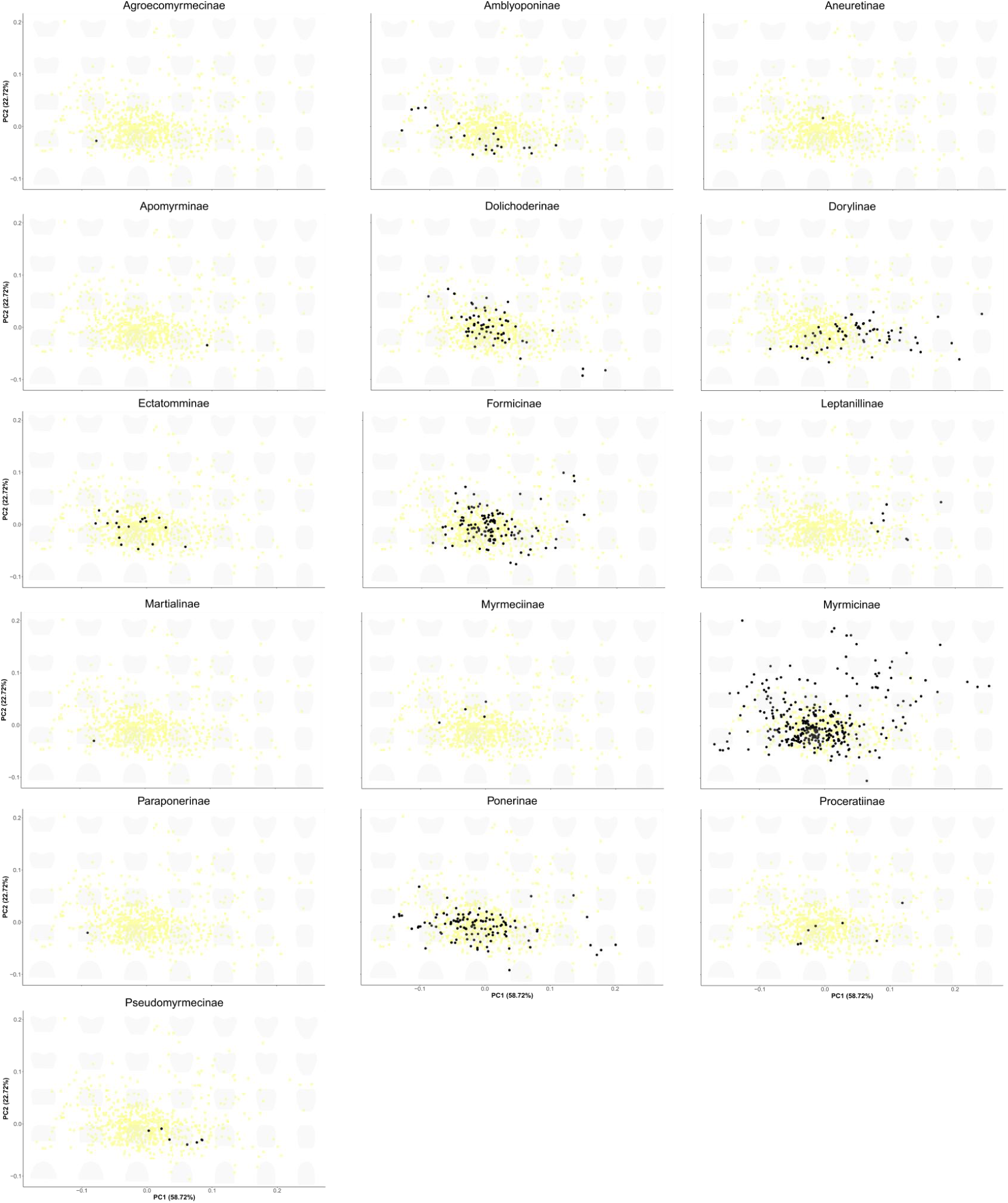
PCA plots representing the morphospace of ant worker 2D head shapes for each subfamily, without allometric effect. In each plot, the black dots represent genera from the subfamily being highlighted, whereas gray dots refer to all remaining genera here considered.

Regarding the phylomorphospace, PC1 and PC2 explained around 58% and 26% of the variation, respectively. Occupation of the phylomorphospace showed, in general, a similar pattern to that observed for the morphospace. The genera *Odontomachus* (Ponerinae) and *Leptomyrmex* (Dolichoderinae) were located at the negative edge of PC2 beyond the range of Myrmicinae (Fig. 3), showing longer and convex heads. Also, *Myopopone* (Amblyoponinae) rivals three Myrmicinae genera in the negative range of PC1, showing a broad and almost convex head (Fig. 3). The expansion of those lineages beyond the range exhibited by Myrmicinae genera is most likely a result of pruning and subsequent averaging of the landmark data.

**Fig.3.**
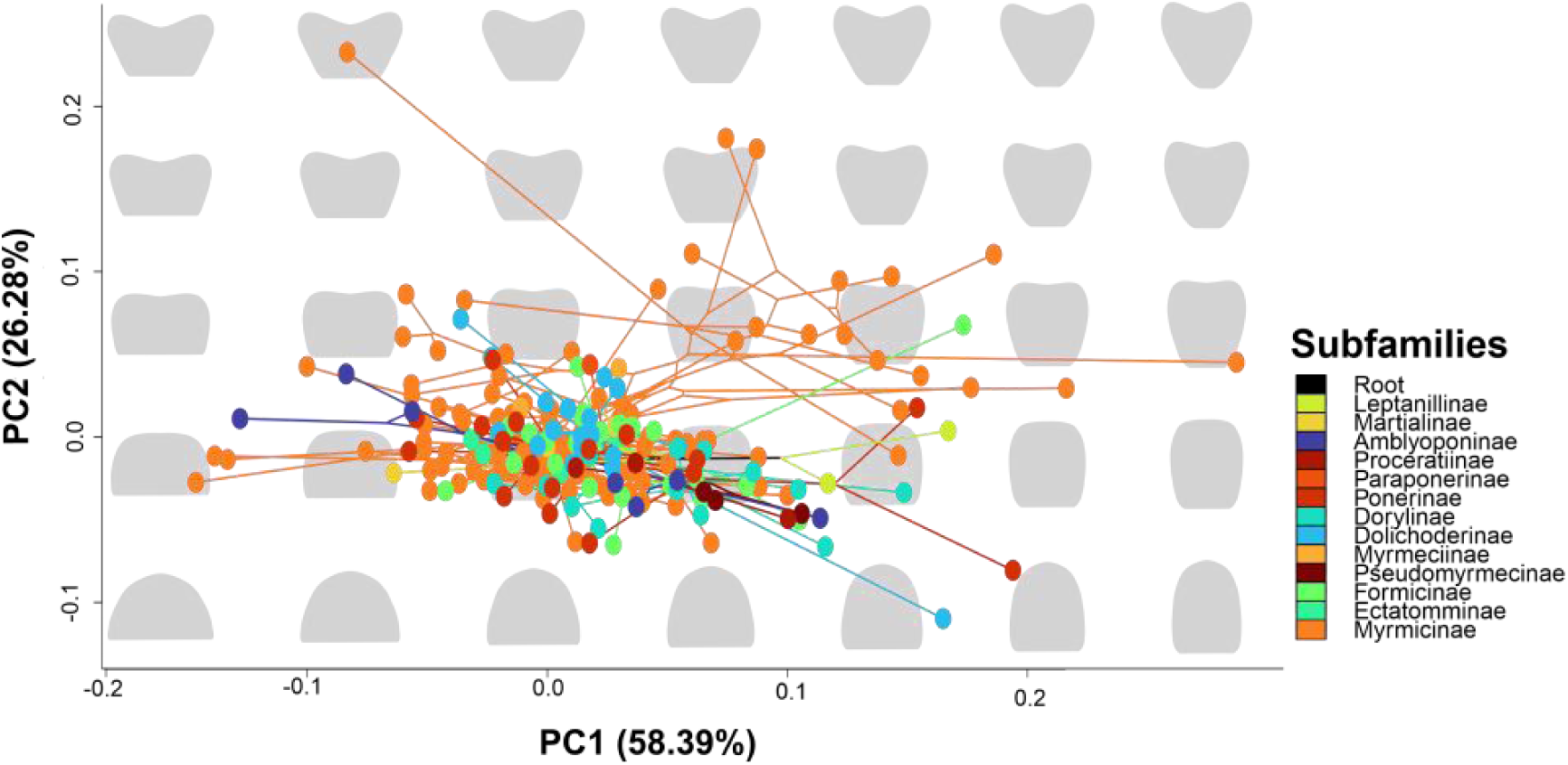
Phylomorphospace of ant worker plane head shapes, without the effect of allometry. Dots represent ant genera and lines depict their phylogenetic relationships, coloured according to subfamily. Idealized plane head shapes are depicted in the background in light grey.

### Finite element analysis

Our simulations revealed trends regarding the morphological variability described in the morphospace. As plane heads becomes wider, stresses concentrate more along their lateral margins and less on their front (Fig.4). Those results indicate that heart-shaped and narrower heads have higher compressive stresses along their sides, which suggests a propensity to spread stresses toward a wider head area than the remaining plane head shapes, which tended to concentrate stresses around the anterior margin of the head (Fig.4). Also, narrower and convex heads, but especially the wider heads with deep vertex depression, need to deal with higher levels of traction stresses around their vertex depression than the remaining head shapes (Fig.4), which could demand some cuticular reinforcement, representing head shapes much less explored in current ant lineages.

**Fig.4.**
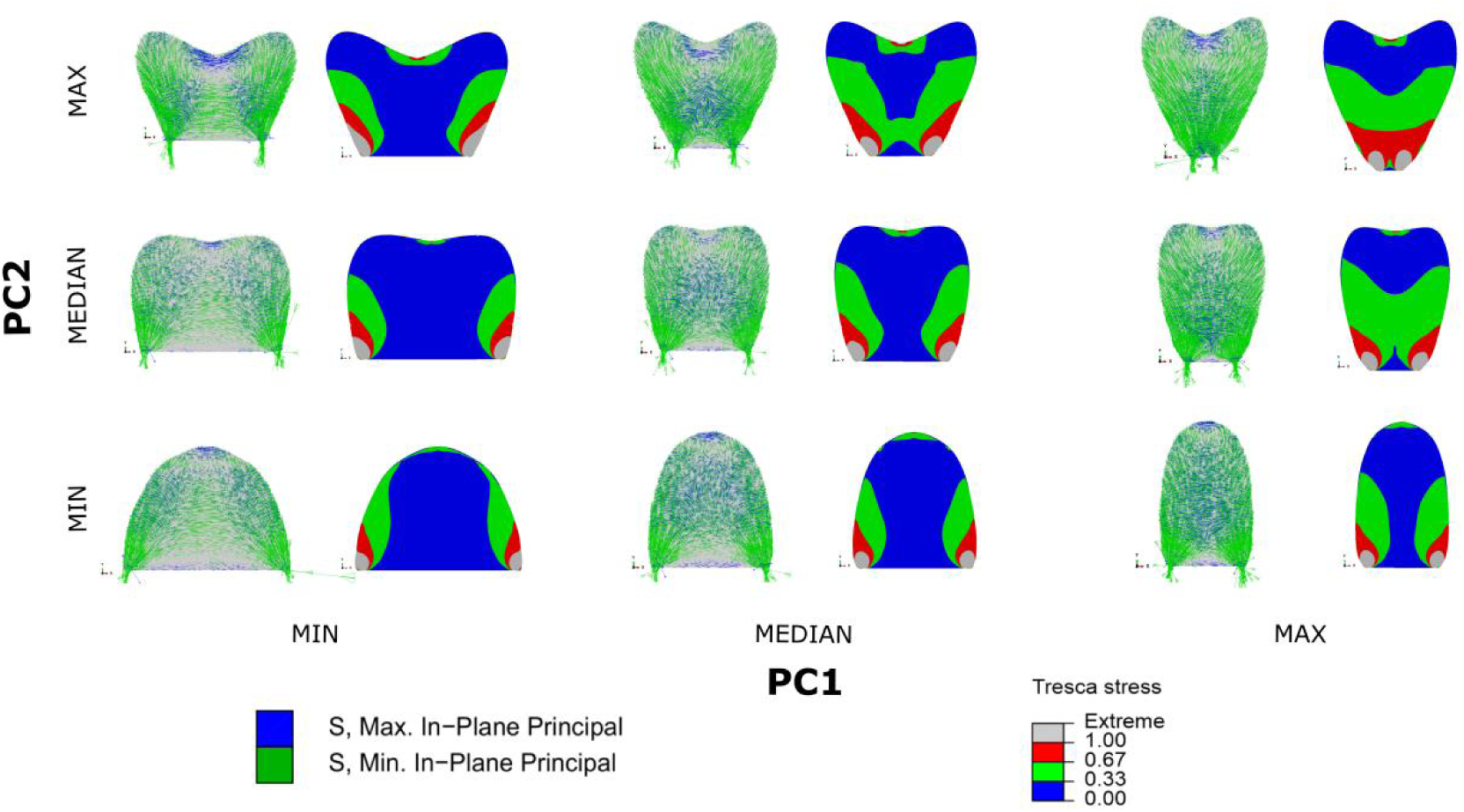
Tensor plots and color maps depicting normalized stress distribution on idealized ant worker head shapes. Tensor plots depict normalized stress magnitude, direction and type (compression or traction) of maximum and minimum stresses, which represent stresses on specific planes were shear stress is zero. Color maps depict normalized Tresca equivalent stress values, without information of stress direction and orientation.

### Intervals method

We performed a PCA following the intervals method with Tresca equivalent stress values to analyze the distribution of non-normalized stress values across head shapes. The first component explained 65% of the variance, being associated with a range of stress values from the lowest (1) towards intermediate (10) stress intervals (Fig.5). Its positive spectrum was mainly affected by the stress distribution of narrow heads with deep vertexal depression located around the positive edges of the phylomorphospace axes (PC1maxPC2max and PC1medianPC2max; Fig.3), and showed a large head area covered with intermediate stresses. Its negative range was more associated with broad heads with shallow vertexal depression (PC1minPC2median) (Fig.5), having a large portion of the head covered with the lowest stress interval. The second PC explained 21% of the variance and was more closely related to the highest stress intervals (Fig.5). The wider and convex head shapes (PC1minPC2min and PC1medianPC2min) were more closely associated with the highest stress intervals, while PC1maxPC2median showed a smaller area of the head covered by those stress ranges, being located at the opposite end of PC2 (Fig.5).

**Fig.5.**
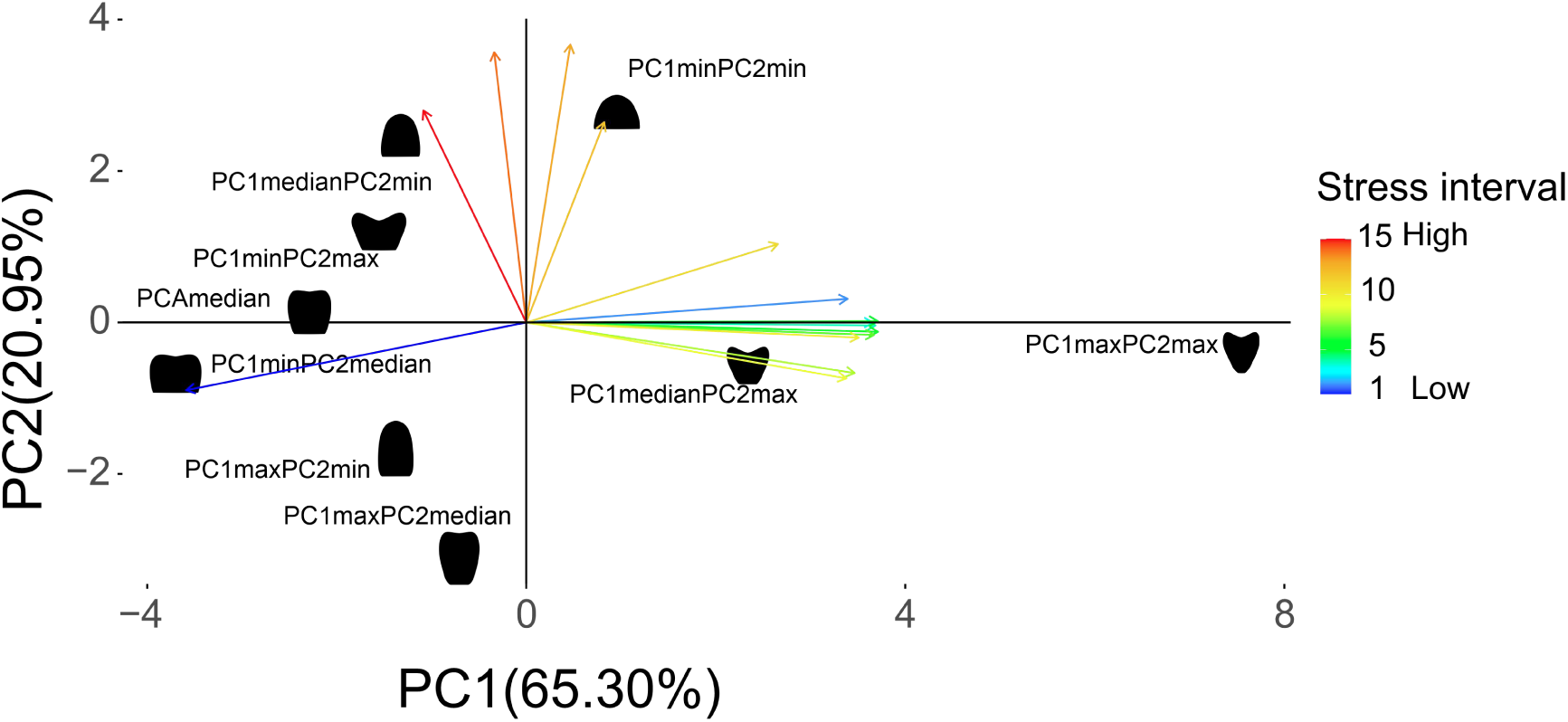
Bivariate plot based on a PCA depicting the distribution of head shapes considering the amount of surface area covered by each of 15 stress intervals, which represent non-normalized and log transformed Tresca equivalent stress values.

## Discussion

Our functional morphology framework contributed to a broad description of ant 2D head shape variation and provided interesting hints about possible biomechanical constraints exhibited by poorly explored head shapes. Regarding worker head morphological diversity, we showed that most current ant lineages have plane head shapes concentrated on a restricted portion of the morphospace, while some lineages, mainly from the subfamily Myrmicinae, more widely explore its limits. This pattern of morphospace occupation indicates that ant head shape evolution is restricted to subtle morphological variations around a common theme. As expected based on observations from *Pheidole* species (Casadei-Ferreira *et al*. 2021, 2022), variation in ant head shape was mainly driven by divergence in head width, length, and degree of vertex depression. Additionally, our results from bite-loading simulations point to potential biomechanical limitations on head shapes placed on the edges of the morphospace and suggest two main avenues of mechanical constraints. One is associated with the significant spread of high-stress levels along the head, as observed on narrow heads with deep vertex depression. The other one regards broader heads whose stresses tend to be notably concentrated around the regions of mandibular articulation, not spreading significantly towards the remaining head area.

The pattern of phylomorphospace occupation recovered here suggests that, despite its documented morphological and ecological variation (Sosiak and Barden 2021), current ant lineages exhibit a generally limited variation in worker head shape. However, they explore the ant head morphospace more widely than stem ants or specialized predators like the so-called hell-ants from the Cretaceous, which occupy distinct regions of the head morphospace but are more restricted than crown ants in their range of distribution (Barden *et al*. 2020). Moreover, we found several instances of morphological convergence at the genus level, even in the less explored regions of the phylomorphospace. For example, at the positive range of PC1 and negative range of PC2, we observed a morphological convergence between a Ponerinae genus (*Odontomachus*) and a Dolichoderinae genus (*Leptomyrmex*). On the opposite end of PC1, three Myrmicinae genera (*Perissomyrmex*, *Pristomyrmex*, and *Dolopomyrmex*) converge in head shape with one Amblyoponinae genus (*Myopopone*). Such a propensity to worker head shape convergence was also noted among *Pheidole* species, regarding lineages distributed on the west and east global hemispheres (Casadei-Ferreira *et al*. 2022), and also among *Cephalotes* species, specifically among workers whose head is adapted to block nest entrances, the soldiers (Powell *et al*. 2020). The limited morphospace exploration and the commonness of convergence suggest that head shape is likely to vary within a restricted range of possibilities.

Another relevant aspect of the phylomorphospace occupation is the specific region that ant genera populated more densely. Since ant workers employ their mandibles to perform most colony tasks (Wilson 1987), it is expected that head shape would reflect an optimization towards a better capacity to store mandibular muscles, especially the adductors, to generate stronger bites. According to several biomechanical and morphological studies (Paul and Gronenberg 1999, Boudinot *et al*. 2021, Püffel *et al*. 2021), such optimization leads to a broader head with a conspicuous vertex depression, which would promote more space for muscle fibre insertion at angles that optimize force generation upon contraction. However, the most common head shape exhibited by current ant species differs from this optimized head, representing a head slightly broader than long, with a shallow vertexal depression. This observation suggests that the provision of space for mandibular muscles may not represent the primary evolutionary pressure driving ant head evolution, being more relevant to some specialized worker castes in a few ant species, like in *Pheidole* spp. and *Atta* spp. major workers (Paul and Gronenberg 1999, Lillico-Ouachour *et al*. 2018, Püffel *et al*. 2021). Another potential shape optimization for muscle arrangement involves the elongation and narrowing of the head, following the development of longer muscle fibres that optimize contraction speed instead of force (Paul and Gronenberg 1999). However, such head shapes are also rarely explored by current ant lineages. Therefore, the phylomorphospace occupation pattern we recovered suggests that an optimization for bite force or speed is not the main driver of ant worker head shape evolution, agreeing with the necessity of ant workers to perform multiple tasks with their mandibles, under distinct mechanical demands (Wilson 1987).

A possible biomechanical explanation for the poor exploration of optimal head shapes for muscle storage is the pattern of stress dissipation they promote. Broader heads tended to accumulate relevant stress levels around the regions of mandibular articulation. A deep vertex depression results in a substantial concentration of tensile stresses on this region. Such stress concentrations on restricted head regions can impose a risk of localised cuticular failure by fatigue, especially when resulting from recurrent biting cycles (Dirks *et al*. 2013), which could be detrimental to the proper functioning of the entire structure, even in those head shapes that were more associated with a larger area of the head covered by the lowest stress intervals (e.g. PC1minPC2median, PC1maxPC2min, and PC1maxPC2median). This is a relevant issue for ant workers who have specialized tasks, like the majors of *Pheidole* and *Atta*, which usually engage in repetitive chewing cycles to process hard food items like insect cuticles, seeds, or leaves, along with in fights for colony defence. Mechanical reinforcements on the most stressed head regions may be necessary to overcome the risks of localised cuticular failure, like the occurrence of cuticular sculptures that locally increase cuticle thickness (Buxton *et al*. 2021) and aid in stopping the failure front, depending on its orientation and microstructure characteristics (Jansen *et al*. 2019). This protective function was recently suggested as a possible explanation for the sculpturing patterns observed on the heads of *Pheidole* major workers (Klunk *et al*. 2023). However, proper mechanical tests are needed to understand the mechanical effects of cuticle sculpturing patterns in this context. Alternatively, a broader head with larger mandibular muscles would also mean a heavier head that shifts the position of the body centre of mass anteriorly (Anderson *et al*. 2020), potentially impairing locomotion, as demonstrated for male stag beetles (Goyens et al. 2015). Accordingly, interspecific variation in ant body center of mass seems to be tightly constrained (Anderson *et al*. 2020). On the other hand, the narrower and longer head shapes that optimize biting speed tend to spread high-stress levels towards a larger area of the head, notably when this shape also shows a deep vertex depression, leading to the accumulation of relevant levels of tensile stresses around this head region. A pattern of wide high-stress spread can thus demand broad and potentially costly cuticular reinforcements, including modifications of the cuticle’s mechanical properties through mechanisms that affect its sclerotization levels (Vincent and Wegst 2004).

The morphospace median head shape, the closest one from the more observed head shape in current ant lineages, seems to exhibit a compromise between the tendency to spread stresses towards the head and the capacity to concentrate stresses on particular head regions. This head shape exhibited a broader coverage of the lowest stress interval than most of the remaining head shapes, and its vertex depression concentrated tensile stresses at proportionally lower levels than those observed on the heads with more depressed posterior margins. Also, the regions of mandibular articulation were proportionally less stressed in this head shape than in most of the remaining idealized heads. Such a combination of mechanical responses to bite loading could help to explain why this morphology is more favoured by current ant lineages, even being morphologically distinct from a head shape optimized to store larger muscles or to improve muscle contraction speed (Paul and Gronenberg 1999, Püffel *et al*. 2021).

### Limitations of our approach in relation to the complexity of ant heads

Planar representations do not provide a satisfactory portrayal of a structure’s mechanical response to loading demands. However, it offers invaluable insights as a first biomechanical approximation (Marcé-Nogué *et al*. 2013), notably when applied to a diverse phylogenetic dataset. Mechanical simulations in planar structures can point to possible sources of weakness, like the depression in the posterior margin of the head or the regions of mandibular articulation, as discussed here for ants. Similar approaches showed that the snout posterior constriction is a delicate region of thalattosuchians (Crocodylomorpha) heads, which might explain why the snout is usually decoupled from the posterior region of the cranium on fossil depositions (Pierce *et al*. 2009). On the same line, heavily stressed zones of *Allosaurus* and *Coelophysis* crania corresponded to skull regions exhibiting thicker bone layers (Rayfield 2005). The weaker zones of trilobite cephalon, which correspond to facial sutures associated with cuticular moulting, were shown to be located in areas of low-stress concentration under burrowing efforts (Esteve *et al*. 2021). However, a necessary step to further explore possible mechanical limitations in ant head shape evolution demands the use of 3D representations of ant heads, preferentially considering its endoskeleton too (Blanke *et al*. 2017, 2018), as well as the potential mechanical effects of the head cuticle sculpturing patterns (Buxton *et al*. 2021). Attempts to investigate the mechanical responses of insect head capsules to bite-loading demonstrated that the varying cuticle thickness and endoskeleton morphology are relevant features for bite-induced stress mitigation (Blanke *et al*. 2017, 2018).

## Conclusions

Biting insects share a general morphology of their biting apparatus, which consists of a pair of mandibles and maxillae, one labium, and the associated musculature that originates from the head capsule. Also, many insects share the dicondylic articulation between the mandibles and head, resulting in a similar pattern of reaction forces generation during bites (Blanke 2019). Therefore, the biting boundary conditions applied here are not unique for ants, being typical for many insects. So, comparable stress patterns can be expected on similar head shapes from other insects, potentially providing insights about head evolution for other insect groups, especially regarding the main tendencies of stress concentration and spread observed with the variation in head dimensions and cuticular depressions.

We provided the first large-scale attempt to quantify the morphological variation of ant worker heads, coupled with a biomechanical investigation that suggested interesting effects between bite-loading demands and head morphological evolution. Based on our morphological quantification, we propose that variation in head width, length, and degree of vertex depression accounts for an expressive amount of variance in the planar head shape of ant lineages, as previously observed in *Pheidole* ants (Casadei-Ferreira *et al*. 2022). Also, according to biomechanical simulations, we point to several potential mechanical limitations exhibited by head shapes from the morphospace edges regarding the intensity and dissipation of stresses. The most commonly explored head shape by current ant lineages exhibited limited stress concentration around the regions of mandibular articulation and vertex depression, allied with mild levels of stress dissipation towards the head, resulting in a broad range of the head covered with lower stresses. In summary, we find support for our hypotheses regarding i) the primary sources of variation in ant workers’ heads; ii) the possibility of biomechanical limitations associated with the rarity of some head shapes among crown ants and; iii) a possible mechanical optimization observed in the more common head shape of current ant lineages.

## Acknowledgments

This study was financed in part by the Coordenação de Aperfeiçoamento de Pessoal de Nível Superior - Brasil (CAPES) - Finance Code 001.

## Author contributions

Cristian L. Klunk, Alexandre Casadei-Ferreira, and Marcio R. Pie conceived the idea of the manuscript and designed the methodology; Cristian L. Klunk and Alexandre Casadei-Ferreira collected the data; Cristian L. Klunk, Alexandre Casadei-Ferreira, and Marco A. Argenta analyzed the data; Cristian L. Klunk and Marcio R. Pie led the writing of the manuscript. All authors contributed critically to the drafts and gave final approval for publication.

