## Supplementary File 1 for "Ant head shape evolved to compromise bite-induced mechanical demands"

**Table S1.** List of species considered for landmark and semilandmark positioning and posterior geometric morphometric analysis. Voucher refers to the image identification on AntWeb online repository (antweb.org).

| Voucher | Subfamily | Genus | Species |
|---|---|---|---|
| antweb1008097 | Dolichoderinae | *Anillidris* | *bruchi* |
| antweb1008361 | Myrmicinae | *Formosimyrma* | *lanyuensis* |
| antweb1008365 | Dolichoderinae | *Bothriomyrmex* | *breviceps* |
| antweb1008367 | Dolichoderinae | *Bothriomyrmex* | *communista* |
| antweb1009013 | Myrmicinae | *Paratopula* | *bauhinia* |
| antweb1038042 | Myrmicinae | *Myrmica* | *incompleta* |
| blf00976(40)-2 | Ponerinae | *Bothroponera* | *cambouei* |
| blf1862(05)-9 | Ponerinae | *Mesoponera* | *ambigua* |
| casent0000078 | Ponerinae | *Dolioponera* | *fustigera* |
| casent0000547 | Dorylinae | *Tanipone* | *hirsuta* |
| casent0002420 | Myrmicinae | *Royidris* | *diminuta* |
| casent0003117 | Myrmicinae | *Metapone* | *emersoni* |
| casent0003125 | Dolichoderinae | *Axinidris* | *gabonica* |
| casent0003155 | Myrmicinae | *Nothomyrmecia* | *macrops* |
| casent0003165 | Paraponerinae | *Paraponera* | *clavata* |
| casent0003196 | Myrmicinae | *Vollenhovia* | *emeryi* |
| casent0003202 | Dorylinae | *Cheliomyrmex* | *morosus* |
| casent0003287 | Myrmicinae | *Pilotrochus* | *besmerus* |
| casent0004352 | Amblyoponinae | *Adetomyrma* | *caputleae* |
| casent0004528 | Myrmicinae | *Metapone* | *madagascarica* |
| casent0004549 | Dorylinae | *Simopone* | *dux* |
| casent0004663 | Ponerinae | *Dinoponera* | *longipes* |
| casent0005175 | Dolichoderinae | *Ochetellus* | *glaber* |
| casent0005317 | Dorylinae | *Syscia* | *augustae* |
| casent0005324 | Dolichoderinae | *Liometopum* | *luctuosum* |
| casent0005359 | Formicinae | *Formica* | *aerata* |
| casent0005465 | Myrmicinae | *Strumigenys* | *abdera* |
| casent0005467 | Myrmicinae | *Strumigenys* | *actis* |
| casent0005697 | Myrmicinae | *Manica* | *bradleyi* |
| casent0005800 | Pseudomyrmecinae | *Pseudomyrmex* | *spinicola* |
| casent0005904 | Agroecomyrmecinae | *Ankylomyrma* | *coronacantha* |
| casent0005922 | Ponerinae | *Centromyrmex* | *bequaerti* |
| casent0006007 | Ponerinae | *Cryptopone* | *gilva* |
| casent0006075 | Myrmicinae | *Harpagoxenus* | *canadensis* |
| casent0006112 | Myrmicinae | *Philidris* | *brunnea* |
| casent0006152 | Myrmicinae | *Liomyrmex* | *gestroi* |
| casent0006786 | Ectatomminae | *Typhlomyrmex* | *pusillus* |
| casent0006819 | Ponerinae | *Platythyrea* | *punctata* |
| casent0007546 | Myrmicinae | *Carebara* | *aberrans* |
| casent0009219 | Dorylinae | *Eciton* | *burchellii* |
| casent0009949 | Dolichoderinae | *Doleromyrma* | *darwiniana* |
| casent0009951 | Dolichoderinae | *Nebothriomyrmex* | *majeri* |
| casent0010788 | Dorylinae | *Cylindromyrmex* | *meinerti* |

| | | | |
|---|---|---|---|
| casent0010791 | Dorylinae | *Nomamyrmex* | *esenbeckii* |
| casent0010799 | Myrmicinae | *Dacetinops* | *concinnus* |
| casent0011746 | Dolichoderinae | *Leptomyrmex* | *erythrocephalus* |
| casent0012024 | Dolichoderinae | *Leptomyrmex* | *darlingtoni* |
| casent0017121 | Myrmicinae | *Royidris* | *admixta* |
| casent0042782 | Myrmicinae | *Vitsika* | *acclivitas* |
| casent0044208 | Myrmicinae | *Malagidris* | *dulcis* |
| casent0047656 | Ponerinae | *Leptogenys* | *acutirostris* |
| casent0052775 | Amblyoponinae | *Prionopelta* | *amabilis* |
| casent0055965 | Dolichoderinae | *Technomyrmex* | *albipes* |
| casent0056913 | Amblyoponinae | *Adetomyrma* | *goblin* |
| casent0058901 | Dolichoderinae | *Technomyrmex* | *anterops* |
| casent0077435 | Myrmicinae | *Eutetramorium* | *mocquerysi* |
| casent0077847 | Formicinae | *Plagiolepis* | *alluaudi* |
| casent0101094 | Myrmicinae | *Melissotarsus* | *insularis* |
| casent0101449 | Dorylinae | *Leptanilloides* | *amazona* |
| casent0101451 | Amblyoponinae | *Fulakora* | *agostii* |
| casent0101453 | Myrmicinae | *Microdaceton* | *exornatum* |
| casent0101591 | Myrmicinae | *Erromyrma* | *latinodis* |
| casent0101696 | Myrmicinae | *Terataner* | *alluaudi* |
| casent0101946 | Myrmicinae | *Chimaeridris* | *burckhardti* |
| casent0102002 | Dorylinae | *Simopone* | *conradti* |
| casent0102086 | Formicinae | *Gesomyrmex* | *chaperi* |
| casent0102121 | Amblyoponinae | *Myopopone* | *castanea* |
| casent0102135 | Ponerinae | *Psalidomyrmex* | *feae* |
| casent0102176 | Amblyoponinae | *Onychomyrmex* | *hedleyi* |
| casent0102193 | Amblyoponinae | *Amblyopone* | *aberrans* |
| casent0102251 | Myrmicinae | *Microdaceton* | *tanyspinosum* |
| casent0102325 | Ponerinae | *Loboponera* | *edentula* |
| casent0102328 | Ponerinae | *Loboponera* | *basalis* |
| casent0102331 | Ponerinae | *Iroponera* | *odax* |
| casent0102362 | Myrmicinae | *Secostruma* | *lethifera* |
| casent0102364 | Myrmicinae | *Lasiomyrma* | *gedensis* |
| casent0102372 | Dolichoderinae | *Froggattella* | *kirbii* |
| casent0102379 | Formicinae | *Bajcaridris* | *theryi* |
| casent0102384 | Formicinae | *Gesomyrmex* | *howardi* |
| casent0102456 | Dolichoderinae | *Ecphorella* | *wellmani* |
| casent0102487 | Amblyoponinae | *Stigmatomma* | *bellii* |
| casent0102512 | Amblyoponinae | *Fulakora* | *armigera* |
| casent0102596 | Myrmicinae | *Strumigenys* | *acheron* |
| casent0102758 | Dolichoderinae | *Liometopum* | *apiculatum* |
| casent0102761 | Dorylinae | *Neivamyrmex* | *agilis* |
| casent0102818 | Formicinae | *Nylanderia* | *austroccidua* |
| casent0102918 | Ponerinae | *Plectroctena* | *anops* |
| casent0102920 | Ponerinae | *Plectroctena* | *cristata* |
| casent0102994 | Ponerinae | *Feroponera* | *ferox* |
| casent0103099 | Myrmicinae | *Perissomyrmex* | *bidentatus* |
| casent0103106 | Formicinae | *Camponotus* | *absquatulator* |
| casent0103121 | Myrmicinae | *Mycetagroicus* | *cerradensis* |
| casent0103128 | Dorylinae | *Acanthostichus* | *arizonensis* |

| | | | |
|---|---|---|---|
| casent0103181 | Formicinae | *Polyrhachis* | *aberrans* |
| casent0103304 | Apomyrminae | *Apomyrma* | *stygia* |
| casent0103385 | Myrmicinae | *Temnothorax* | *nevadensis* |
| casent0103395 | Myrmicinae | *Patagonomyrmex* | *odoratus* |
| casent0103494 | Myrmicinae | *Formicoxenus* | *hirticornis* |
| casent0103517 | Myrmicinae | *Cryptomyrmex* | *longinodus* |
| casent0103673 | Formicinae | *Camponotus* | *floridanus* |
| casent0103752 | Myrmicinae | *Cardiocondyla* | *obscurior* |
| casent0104060 | Myrmicinae | *Mycetosoritis* | *hartmanni* |
| casent0104325 | Myrmicinae | *Indomyrma* | *dasypyx* |
| casent0104326 | Myrmicinae | *Amoimyrmex* | *striatus* |
| casent0104575 | Amblyoponinae | *Amblyopone* | *australis* |
| casent0104586 | Ponerinae | *Emeryopone* | *buttelreepeni* |
| casent0104609 | Myrmicinae | *Malagidris* | *belti* |
| casent0104667 | Dorylinae | *Leptanilloides* | *anae* |
| casent0104676 | Dorylinae | *Leptanilloides* | *biconstricta* |
| casent0104767 | Formicinae | *Formica* | *adamsi* |
| casent0104822 | Myrmicinae | *Formicoxenus* | *quebecensis* |
| casent0104847 | Myrmicinae | *Leptothorax* | *muscorum* |
| casent0104904 | Dolichoderinae | *Linepithema* | *anathema* |
| casent0104984 | Dorylinae | *Yunodorylus* | *sexspinus* |
| casent0105632 | Myrmicinae | *Veromessor* | *lobognathus* |
| casent0105862 | Myrmicinae | *Temnothorax* | *adustus* |
| casent0106006 | Formicinae | *Calomyrmex* | *albertisi* |
| casent0106082 | Ectatomminae | *Typhlomyrmex* | *rogenhoferi* |
| casent0106119 | Dolichoderinae | *Linepithema* | *humile* |
| casent0106177 | Myrmicinae | *Tyrannomyrmex* | *legatus* |
| casent0106181 | Martialinae | *Martialis* | *heureka* |
| casent0106204 | Dorylinae | *Parasyscia* | *desposyne* |
| casent0106206 | Myrmicinae | *Tranopelta* | *gilva* |
| casent0106211 | Myrmicinae | *Austromorium* | *flavigaster* |
| casent0106220 | Myrmicinae | *Dacetinops* | *cibdelus* |
| casent0106225 | Myrmicinae | *Lophomyrmex* | *ambiguus* |
| casent0106227 | Dorylinae | *Labidus* | *praedator* |
| casent0106233 | Dorylinae | *Cerapachys* | *jacobsoni* |
| casent0106243 | Myrmicinae | *Acanthomyrmex* | *ferox* |
| casent0106263 | Myrmicinae | *Trichomyrmex* | *criniceps* |
| casent0106343 | Myrmicinae | *Rostromyrmex* | *pasohensis* |
| casent0125111 | Formicinae | *Anoplolepis* | *gracilipes* |
| casent0127094 | Dolichoderinae | *Leptomyrmex* | *cnemidatus* |
| casent0129291 | Formicinae | *Prenolepis* | *fustinoda* |
| casent0129840 | Amblyoponinae | *Mystrium* | *barrybressleri* |
| casent0130146 | Dolichoderinae | *Aptinoma* | *mangabe* |
| casent0137608 | Myrmicinae | *Nesomyrmex* | *angulatus* |
| casent0145852 | Myrmicinae | *Eutetramorium* | *monticellii* |
| casent0150743 | Dorylinae | *Lividopone* | *livida* |
| casent0170535 | Formicinae | *Anoplolepis* | *custodiens* |
| casent0170538 | Formicinae | *Acropyga* | *arnoldi* |
| casent0170958 | Myrmicinae | *Malagidris* | *alperti* |
| casent0171177 | Myrmicinae | *Propodilobus* | *pingorum* |

| | | | |
|---|---|---|---|
| casent0171178 | Myrmicinae | *Lordomyrma* | *bhutanensis* |
| casent0172082 | Amblyoponinae | *Mystrium* | *camillae* |
| casent0172109 | Dorylinae | *Zasphinctus* | *caledonicus* |
| casent0172203 | Amblyoponinae | *Onychomyrmex* | *glauerti* |
| casent0172264 | Myrmicinae | *Peronomyrmex* | *greavesi* |
| casent0172278 | Myrmicinae | *Poecilomyrma* | *senirewae* |
| casent0172284 | Myrmicinae | *Huberia* | *striata* |
| casent0172291 | Dolichoderinae | *Loweriella* | *boltoni* |
| casent0172294 | Myrmicinae | *Huberia* | *brounii* |
| casent0172297 | Amblyoponinae | *Onychomyrmex* | *doddi* |
| casent0172316 | Myrmicinae | *Peronomyrmex* | *overbecki* |
| casent0172339 | Ponerinae | *Austroponera* | *castaneicolor* |
| casent0172360 | Myrmicinae | *Orectognathus* | *antennatus* |
| casent0172424 | Ponerinae | *Ponera* | *augusta* |
| casent0172425 | Ponerinae | *Ponera* | *clavicornis* |
| casent0172430 | Ponerinae | *Paltothyreus* | *tarsatus* |
| casent0172431 | Ponerinae | *Buniapone* | *amblyops* |
| casent0172441 | Myrmicinae | *Mayriella* | *ebbei* |
| casent0172443 | Myrmicinae | *Mayriella* | *sharpi* |
| casent0172451 | Myrmicinae | *Dacetinops* | *cirrosus* |
| casent0172464 | Myrmicinae | *Eurhopalothrix* | *australis* |
| casent0172466 | Myrmicinae | *Mesostruma* | *bella* |
| casent0172468 | Myrmicinae | *Mesostruma* | *eccentrica* |
| casent0172783 | Myrmicinae | *Gauromyrmex* | *bengkalisi* |
| casent0172786 | Myrmicinae | *Gauromyrmex* | *acanthinus* |
| casent0172807 | Amblyoponinae | *Fulakora* | *orizabana* |
| casent0172816 | Amblyoponinae | *Amblyopone* | *clarki* |
| casent0172824 | Myrmicinae | *Chimaeridris* | *boltoni* |
| casent0172947 | Formicinae | *Prolasius* | *advena* |
| casent0173029 | Ponerinae | *Thaumatomyrmex* | *cochlearis* |
| casent0173051 | Dorylinae | *Zasphinctus* | *asper* |
| casent0173056 | Dorylinae | *Eusphinctus* | *furcatus* |
| casent0173159 | Myrmicinae | *Formicoxenus* | *nitidulus* |
| casent0173374 | Myrmicinae | *Patagonomyrmex* | *angustus* |
| casent0173381 | Ponerinae | *Dinoponera* | *australis* |
| casent0173474 | Formicinae | *Brachymyrmex* | *aphidicola* |
| casent0173500 | Dorylinae | *Acanthostichus* | *brevicornis* |
| casent0173502 | Dorylinae | *Cylindromyrmex* | *brasiliensis* |
| casent0173514 | Dorylinae | *Labidus* | *mars* |
| casent0173533 | Ponerinae | *Anochetus* | *altisquamis* |
| casent0173538 | Ponerinae | *Odontomachus* | *meinerti* |
| casent0173540 | Ectatomminae | *Acanthoponera* | *mucronata* |
| casent0173546 | Myrmicinae | *Eutetramorium* | *parvum* |
| casent0173595 | Dorylinae | *Labidus* | *coecus* |
| casent0173637 | Ponerinae | *Streblognathus* | *peetersi* |
| casent0173652 | Myrmicinae | *Trachymyrmex* | *desertorum* |
| casent0173838 | Dolichoderinae | *Dolichoderus* | *lamellosus* |
| casent0173879 | Myrmicinae | *Lachnomyrmex* | *lattkei* |
| casent0173880 | Myrmicinae | *Lachnomyrmex* | *fernandezi* |
| casent0173906 | Formicinae | *Cladomyrma* | *hewitti* |

| | | | |
|---|---|---|---|
| casent0173913 | Formicinae | *Cladomyrma* | *andrei* |
| casent0173917 | Formicinae | *Cladomyrma* | *crypteroniae* |
| casent0173968 | Myrmicinae | *Hylomyrma* | *balzani* |
| casent0173986 | Myrmicinae | *Monomorium* | *pharaonis* |
| casent0173988 | Myrmicinae | *Mycocepurus* | *goeldii* |
| casent0173990 | Myrmicinae | *Myrmicocrypta* | *squamosa* |
| casent0178018 | Myrmicinae | *Pheidole* | *fimbriata* |
| casent0178023 | Dolichoderinae | *Arnoldius* | *flavus* |
| casent0178098 | Myrmicinae | *Oxyepoecus* | *bruchi* |
| casent0178105 | Myrmicinae | *Mycetomoellerius* | *dichrous* |
| casent0178167 | Myrmicinae | *Rogeria* | *alzatei* |
| casent0178203 | Ponerinae | *Phrynoponera* | *pulchella* |
| casent0178228 | Ponerinae | *Phrynoponera* | *gabonensis* |
| casent0178237 | Myrmicinae | *Diaphoromyrma* | *sofiae* |
| casent0178273 | Dolichoderinae | *Technomyrmex* | *andrei* |
| casent0178287 | Myrmicinae | *Calyptomyrmex* | *brunneus* |
| casent0178293 | Myrmicinae | *Cyphoidris* | *spinosa* |
| casent0178294 | Myrmicinae | *Melissotarsus* | *weissi* |
| casent0178295 | Myrmicinae | *Microdaceton* | *tibialis* |
| casent0178340 | Ponerinae | *Centromyrmex* | *alfaroi* |
| casent0178347 | Leptanillinae | *Opamyrma* | *hungvuong* |
| casent0178387 | Myrmicinae | *Syllophopsis* | *australica* |
| casent0178452 | Ponerinae | *Brachyponera* | *atrata* |
| casent0178489 | Myrmicinae | *Daceton* | *armigerum* |
| casent0178493 | Formicinae | *Agraulomyrmex* | *meridionalis* |
| casent0178496 | Formicinae | *Alloformica* | *nitidior* |
| casent0178502 | Formicinae | *Overbeckia* | *subclavata* |
| casent0178514 | Formicinae | *Rossomyrmex* | *proformicarum* |
| casent0178522 | Myrmicinae | *Recurvidris* | *hebe* |
| casent0178524 | Myrmicinae | *Romblonella* | *palauensis* |
| casent0178526 | Myrmicinae | *Proatta* | *butteli* |
| casent0178537 | Myrmicinae | *Kartidris* | *matertera* |
| casent0178543 | Myrmicinae | *Lophomyrmex* | *birmanus* |
| casent0178565 | Myrmicinae | *Dilobocondyla* | *fouqueti* |
| casent0178567 | Myrmicinae | *Dolopomyrmex* | *pilatus* |
| casent0178575 | Myrmicinae | *Acanthomyrmex* | *basispinosus* |
| casent0178593 | Myrmicinae | *Stereomyrmex* | *horni* |
| casent0178635 | Myrmicinae | *Paratrachymyrmex* | *cornetzi* |
| casent0178653 | Myrmicinae | *Rogeria* | *belti* |
| casent0178703 | Ponerinae | *Thaumatomyrmex* | *atrox* |
| casent0178718 | Myrmicinae | *Acanthognathus* | *ocellatus* |
| casent0178733 | Myrmicinae | *Talaridris* | *mandibularis* |
| casent0178751 | Ponerinae | *Promyopias* | *silvestrii* |
| casent0178759 | Formicinae | *Paraparatrechina* | *albipes* |
| casent0178772 | Myrmicinae | *Harpagoxenus* | *sublaevis* |
| casent0178839 | Myrmicinae | *Goniomma* | *hispanicum* |
| casent0178842 | Myrmicinae | *Rhopalomastix* | *johorensis* |
| casent0178849 | Dorylinae | *Sphinctomyrmex* | *schoerederi* |
| casent0178850 | Dorylinae | *Sphinctomyrmex* | *stali* |
| casent0179461 | Dorylinae | *Chrysapace* | *jacobsoni* |

| | | | |
|---|---|---|---|
| casent0179468 | Myrmicinae | *Paratrachymyrmex* | *bugnioni* |
| casent0179559 | Formicinae | *Polyergus* | *breviceps* |
| casent0179594 | Myrmicinae | *Ochetomyrmex* | *neopolitus* |
| casent0179897 | Formicinae | *Lasius* | *niger* |
| casent0179982 | Ectatomminae | *Gnamptogenys* | *alfaroi* |
| casent0205996 | Amblyoponinae | *Adetomyrma* | *bressleri* |
| casent0217032 | Leptanillinae | *Anomalomyrma* | *boltoni* |
| casent0217049 | Formicinae | *Aphomomyrmex* | *afer* |
| casent0217050 | Myrmicinae | *Kempfidris* | *inusualis* |
| casent0217130 | Dolichoderinae | *Axinidris* | *bidens* |
| casent0217157 | Dorylinae | *Dorylus* | *affinis* |
| casent0217159 | Dorylinae | *Dorylus* | *braunsi* |
| casent0217200 | Formicinae | *Plagiolepis* | *boltoni* |
| casent0217258 | Myrmicinae | *Patagonomyrmex* | *laevigatus* |
| casent0217322 | Amblyoponinae | *Xymmer* | *muticus* |
| casent0217418 | Formicinae | *Polyrhachis* | *abdita* |
| casent0217470 | Dorylinae | *Eciton* | *hamatum* |
| casent0217516 | Ponerinae | *Diacamma* | *rugosum* |
| casent0217536 | Ponerinae | *Odontomachus* | *affinis* |
| casent0217550 | Ponerinae | *Mayaponera* | *arhuaca* |
| casent0217553 | Ponerinae | *Pseudoponera* | *cognata* |
| casent0217812 | Formicinae | *Stigmacros* | *aemula* |
| casent0217815 | Myrmicinae | *Amoimyrmex* | *silvestrii* |
| casent0217816 | Myrmicinae | *Novomessor* | *ensifer* |
| casent0217908 | Myrmicinae | *Podomyrma* | *adelaidae* |
| casent0219919 | Dolichoderinae | *Anonychomyrma* | *dimorpha* |
| casent0220221 | Leptanillinae | *Anomalomyrma* | *helenae* |
| casent0221916 | Myrmicinae | *Aphaenogaster* | *occidentalis* |
| casent0235147 | Myrmicinae | *Anillomyrma* | *decamera* |
| casent0235148 | Myrmicinae | *Dacatria* | *templaris* |
| casent0235329 | Myrmicinae | *Pogonomyrmex* | *barbatus* |
| casent0235341 | Leptanillinae | *Protanilla* | *bicolor* |
| casent0235373 | Myrmicinae | *Calyptomyrmex* | *duhun* |
| casent0235475 | Proceratiinae | *Discothyrea* | *aisnetu* |
| casent0235532 | Myrmicinae | *Monomorium* | *affabile* |
| casent0235605 | Ponerinae | *Euponera* | *brunoi* |
| casent0235607 | Ponerinae | *Megaponera* | *analis* |
| casent0235609 | Ponerinae | *Parvaponera* | *suspecta* |
| casent0235681 | Amblyoponinae | *Prionopelta* | *aethiopica* |
| casent0235739 | Myrmicinae | *Orectognathus* | *bottegoi* |
| casent0235778 | Myrmicinae | *Tetramorium* | *aculeatum* |
| casent0246060 | Myrmicinae | *Meranoplus* | *bicolor* |
| casent0246694 | Ectatomminae | *Heteroponera* | *dentinodis* |
| casent0247296 | Myrmicinae | *Tetramorium* | *adamsi* |
| casent0249114 | Dorylinae | *Simopone* | *gressitti* |
| casent0249125 | Ponerinae | *Odontoponera* | *denticulata* |
| casent0249149 | Ponerinae | *Pachycondyla* | *harpax* |
| casent0249163 | Ponerinae | *Neoponera* | *unidentata* |
| casent0249173 | Ponerinae | *Ectomomyrmex* | *acutus* |
| casent0249178 | Ponerinae | *Austroponera* | *rufonigra* |

| | | | |
|---|---|---|---|
| casent0249185 | Ponerinae | *Pseudoneoponera* | *excavata* |
| casent0249204 | Ponerinae | *Hagensia* | *peringueyi* |
| casent0249241 | Ponerinae | *Psalidomyrmex* | *foveolatus* |
| casent0249249 | Ponerinae | *Streblognathus* | *aethiopicus* |
| casent0249261 | Proceratiinae | *Proceratium* | *australe* |
| casent0249288 | Dorylinae | *Lioponera* | *aberrans* |
| casent0249379 | Formicinae | *Colobopsis* | *abdita* |
| casent0249455 | Dorylinae | *Eciton* | *dulcium* |
| casent0249516 | Dolichoderinae | *Anonychomyrma* | *biconvexa* |
| casent0249635 | Formicinae | *Paratrechina* | *kohli* |
| casent0249737 | Dolichoderinae | *Linepithema* | *angulatum* |
| casent0249757 | Dolichoderinae | *Philidris* | *cordata* |
| casent0249770 | Dolichoderinae | *Tapinoma* | *sessile* |
| casent0249807 | Dolichoderinae | *Turneria* | *arbusta* |
| casent0249892 | Formicinae | *Cataglyphis* | *abyssinica* |
| casent0249906 | Myrmicinae | *Trichomyrmex* | *abyssinicus* |
| casent0249914 | Formicinae | *Acropyga* | *ambigua* |
| casent0250833 | Myrmicinae | *Ocymyrmex* | *alacer* |
| casent0257669 | Ponerinae | *Mesoponera* | *caffraria* |
| casent0260421 | Ponerinae | *Dinoponera* | *gigantea* |
| casent0260430 | Ponerinae | *Hypoponera* | *nitidula* |
| casent0260462 | Amblyoponinae | *Prionopelta* | *amieti* |
| casent0260514 | Ponerinae | *Belonopelta* | *deletrix* |
| casent0260518 | Ponerinae | *Cryptopone* | *fusciceps* |
| casent0270204 | Myrmicinae | *Carebara* | *affinis* |
| casent0280278 | Formicinae | *Dinomyrmex* | *gigas* |
| casent0280334 | Formicinae | *Echinopla* | *australis* |
| casent0280428 | Formicinae | *Lasiophanes* | *atriventris* |
| casent0280453 | Formicinae | *Lasius* | *alienoflavus* |
| casent0280493 | Formicinae | *Melophorus* | *ankylochaetes* |
| casent0280518 | Formicinae | *Myrmecocystus* | *depilis* |
| casent0280537 | Formicinae | *Myrmecorhynchus* | *emeryi* |
| casent0280560 | Formicinae | *Notoncus* | *hickmani* |
| casent0280568 | Formicinae | *Notostigma* | *foreli* |
| casent0280570 | Formicinae | *Notostigma* | *carazzii* |
| casent0280684 | Myrmicinae | *Acanthognathus* | *brevicornis* |
| casent0280780 | Myrmicinae | *Rhopalothrix* | *isthmica* |
| casent0280937 | Myrmicinae | *Ocymyrmex* | *cavatodorsatus* |
| casent0280972 | Dorylinae | *Aenictus* | *arabicus* |
| casent0281044 | Formicinae | *Polyergus* | *longicornis* |
| casent0281138 | Formicinae | *Oecophylla* | *smaragdina* |
| casent0281139 | Formicinae | *Opisthopsis* | *diademata* |
| casent0281158 | Formicinae | *Petalomyrmex* | *phylax* |
| casent0281306 | Ectatomminae | *Rhytidoponera* | *acanthoponeroides* |
| casent0281337 | Ectatomminae | *Rhytidoponera* | *aenescens* |
| casent0281460 | Formicinae | *Zatania* | *albimaculata* |
| casent0281461 | Formicinae | *Zatania* | *gibberosa* |
| casent0281462 | Formicinae | *Prenolepis* | *fisheri* |
| casent0281489 | Formicinae | *Pseudonotoncus* | *hirsutus* |
| casent0281511 | Ectatomminae | *Gnamptogenys* | *acuminata* |

| | | | |
|---|---|---|---|
| casent0281762 | Myrmicinae | *Mycetophylax* | *bigibbosus* |
| casent0281796 | Myrmicinae | *Pristomyrmex* | *bicolor* |
| casent0281863 | Proceratiinae | *Discothyrea* | *clavicornis* |
| casent0281874 | Pseudomyrmecinae | *Tetraponera* | *nodosa* |
| casent0281878 | Ponerinae | *Anochetus* | *agilis* |
| casent0281904 | Ponerinae | *Neoponera* | *agilis* |
| casent0281906 | Ponerinae | *Myopias* | *breviloba* |
| casent0281913 | Ponerinae | *Hypoponera* | *aliena* |
| casent0281979 | Dorylinae | *Neocerapachys* | *neotropicus* |
| casent0281981 | Dorylinae | *Eusphinctus* | *taylori* |
| casent0289205 | Ponerinae | *Fisheropone* | *ambigua* |
| casent0317395 | Amblyoponinae | *Mystrium* | *eques* |
| casent0317589 | Ponerinae | *Euponera* | *agnivo* |
| casent0317590 | Ponerinae | *Euponera* | *antsiraka* |
| casent0374638 | Ponerinae | *Mayaponera* | *becculata* |
| casent0401720 | Myrmicinae | *Dicroaspis* | *cryptocera* |
| casent0401737 | Ponerinae | *Boloponera* | *vicans* |
| casent0405993 | Myrmicinae | *Cyphoidris* | *exalta* |
| casent0406793 | Ponerinae | *Asphinctopone* | *silvestrii* |
| casent0406851 | Myrmicinae | *Baracidris* | *sitra* |
| casent0407980 | Formicinae | *Nylanderia* | *amblyops* |
| casent0410479 | Dorylinae | *Tanipone* | *pilosa* |
| casent0417143 | Ponerinae | *Asphinctopone* | *differens* |
| casent0417147 | Ponerinae | *Centromyrmex* | *angolensis* |
| casent0417505 | Ponerinae | *Phrynoponera* | *bequaerti* |
| casent0418269 | Dolichoderinae | *Aptinoma* | *antongil* |
| casent0426618 | Myrmicinae | *Melissotarsus* | *emeryi* |
| casent0428251 | Myrmicinae | *Vitsika* | *crebra* |
| casent0434565 | Myrmicinae | *Ancyridris* | *polyrhachioides* |
| casent0439855 | Dolichoderinae | *Ravavy* | *miafina* |
| casent0453836 | Myrmicinae | *Royidris* | *clarinodis* |
| casent0454372 | Formicinae | *Paratrechina* | *ankarana* |
| casent0491610 | Myrmicinae | *Syllophopsis* | *fisheri* |
| casent0492142 | Dorylinae | *Parasyscia* | *imerinensis* |
| casent0492248 | Dorylinae | *Tanipone* | *aglandula* |
| casent0494750 | Myrmicinae | *Vitsika* | *brevis* |
| casent0603528 | Proceratiinae | *Proceratium* | *mancum* |
| casent0604602 | Myrmicinae | *Stenamma* | *andersoni* |
| casent0612214 | Myrmicinae | *Apterostigma* | *dentigerum* |
| casent0613604 | Myrmicinae | *Apterostigma* | *auriculatum* |
| casent0615273 | Formicinae | *Brachymyrmex* | *bicolor* |
| casent0616568 | Myrmicinae | *Epelysidris* | *brocha* |
| casent0617007 | Myrmicinae | *Romblonella* | *longinoi* |
| casent0618628 | Formicinae | *Zatania* | *gloriosa* |
| casent0619175 | Dorylinae | *Neivamyrmex* | *adnepos* |
| casent0619176 | Dorylinae | *Neivamyrmex* | *alfaroi* |
| casent0627381 | Myrmicinae | *Octostruma* | *convallis* |
| casent0627826 | Myrmicinae | *Adelomyrmex* | *marginodus* |
| casent0629582 | Myrmicinae | *Rhopalothrix* | *andersoni* |
| casent0629589 | Myrmicinae | *Rhopalothrix* | *apertor* |

| | | | |
|---|---|---|---|
| casent0630965 | Myrmicinae | *Megalomyrmex* | *adamsae* |
| casent0635091 | Ponerinae | *Rasopone* | *cubitalis* |
| casent0635430 | Myrmicinae | *Pheidole* | *absurda* |
| casent0635809 | Ponerinae | *Rasopone* | *costaricensis* |
| casent0637306 | Myrmicinae | *Hylomyrma* | *montana* |
| casent0644416 | Dorylinae | *Syscia* | *tolteca* |
| casent0646942 | Myrmicinae | *Solenopsis* | *invicta* |
| casent0840860 | Myrmicinae | *Syllophopsis* | *aureorugosa* |
| casent0844332 | Myrmicinae | *Veromessor* | *chamberlini* |
| casent0900003 | Myrmicinae | *Mesostruma* | *browni* |
| casent0900004 | Myrmicinae | *Colobostruma* | *biconvexa* |
| casent0900010 | Myrmicinae | *Colobostruma* | *australis* |
| casent0900022 | Myrmicinae | *Epopostruma* | *alata* |
| casent0900023 | Myrmicinae | *Epopostruma* | *angela* |
| casent0900327 | Myrmicinae | *Atopomyrmex* | *calpocalycola* |
| casent0900500 | Myrmicinae | *Myrmicaria* | *arachnoides* |
| casent0900515 | Ectatomminae | *Ectatomma* | *brunneum* |
| casent0900557 | Ectatomminae | *Gnamptogenys* | *annulata* |
| casent0900926 | Myrmicinae | *Octostruma* | *balzani* |
| casent0900928 | Myrmicinae | *Eurhopalothrix* | *gravis* |
| casent0900943 | Myrmicinae | *Protalaridris* | *armata* |
| casent0900951 | Ponerinae | *Ophthalmopone* | *berthoudi* |
| casent0900952 | Myrmicinae | *Rogeria* | *blanda* |
| casent0900983 | Myrmicinae | *Tetheamyrma* | *subspongia* |
| casent0900990 | Myrmicinae | *Calyptomyrmex* | *shasu* |
| casent0901013 | Myrmicinae | *Baracidris* | *meketra* |
| casent0901014 | Myrmicinae | *Baracidris* | *pilosa* |
| casent0901020 | Myrmicinae | *Cryptomyrmex* | *boltoni* |
| casent0901021 | Myrmicinae | *Tyrannomyrmex* | *rex* |
| casent0901074 | Myrmicinae | *Tetramorium* | *adelphon* |
| casent0901255 | Myrmicinae | *Kartidris* | *nyos* |
| casent0901256 | Myrmicinae | *Kartidris* | *galos* |
| casent0901261 | Myrmicinae | *Lenomyrmex* | *wardi* |
| casent0901347 | Dorylinae | *Cerapachys* | *antennatus* |
| casent0901605 | Myrmicinae | *Pheidole* | *aberrans* |
| casent0901663 | Myrmicinae | *Blepharidatta* | *brasiliensis* |
| casent0901700 | Myrmicinae | *Acanthomyrmex* | *concavus* |
| casent0901741 | Myrmicinae | *Paratopula* | *ankistra* |
| casent0901743 | Myrmicinae | *Paratopula* | *catocha* |
| casent0901769 | Myrmicinae | *Leptothorax* | *acervorum* |
| casent0901824 | Formicinae | *Polyrhachis* | *abdominalis* |
| casent0901921 | Dolichoderinae | *Philidris* | *cruda* |
| casent0901992 | Myrmicinae | *Vombisidris* | *nahet* |
| casent0901994 | Myrmicinae | *Rotastruma* | *stenoceps* |
| casent0901995 | Myrmicinae | *Rotastruma* | *recava* |
| casent0901996 | Myrmicinae | *Romblonella* | *elysii* |
| casent0902000 | Myrmicinae | *Podomyrma* | *alae* |
| casent0902054 | Myrmicinae | *Recurvidris* | *browni* |
| casent0902055 | Myrmicinae | *Recurvidris* | *pickburni* |
| casent0902174 | Myrmicinae | *Anillomyrma* | *tridens* |

| | | | |
|---|---|---|---|
| casent0902176 | Myrmicinae | *Allomerus* | *decemarticulatus* |
| casent0902177 | Myrmicinae | *Allomerus* | *septemarticulatus* |
| casent0902288 | Myrmicinae | *Chelaner* | *albipes* |
| casent0902332 | Myrmicinae | *Megalomyrmex* | *bidentatus* |
| casent0902411 | Ectatomminae | *Heteroponera* | *brounii* |
| casent0902494 | Ponerinae | *Brachyponera* | *chinensis* |
| casent0902502 | Ponerinae | *Pseudoneoponera* | *denticulata* |
| casent0902520 | Ponerinae | *Myopias* | *castaneicola* |
| casent0902668 | Dorylinae | *Nomamyrmex* | *hartigii* |
| casent0902681 | Dorylinae | *Aenictus* | *alticola* |
| casent0902700 | Dorylinae | *Vicinopone* | *conciliatrix* |
| casent0902716 | Dorylinae | *Yunodorylus* | *paradoxus* |
| casent0902717 | Dorylinae | *Yunodorylus* | *doryloides* |
| CASENT0902774 | Leptanillinae | *Leptanilla* | *buddhista* |
| casent0902804 | Myrmeciinae | *Myrmecia* | *varians* |
| casent0902837 | Pseudomyrmecinae | *Myrcidris* | *epicharis* |
| casent0902845 | Pseudomyrmecinae | *Pseudomyrmex* | *gracilis* |
| casent0902927 | Dolichoderinae | *Turneria* | *bidentata* |
| casent0902931 | Dolichoderinae | *Axinidris* | *ghanensis* |
| casent0902944 | Dolichoderinae | *Dolichoderus* | *abruptus* |
| casent0902997 | Dolichoderinae | *Papyrius* | *nitidus* |
| casent0902998 | Dolichoderinae | *Gracilidris* | *pombero* |
| casent0903002 | Dolichoderinae | *Azteca* | *alfari* |
| casent0903092 | Dolichoderinae | *Iridomyrmex* | *alpinus* |
| casent0903110 | Formicinae | *Tapinolepis* | *deceptor* |
| casent0903113 | Formicinae | *Myrmelachista* | *ambigua* |
| casent0903172 | Formicinae | *Acropyga* | *acutiventris* |
| casent0903195 | Formicinae | *Myrmecocystus* | *colei* |
| casent0903196 | Formicinae | *Myrmecocystus* | *creightoni* |
| casent0903205 | Formicinae | *Lasius* | *sitiens* |
| casent0903229 | Formicinae | *Prolasius* | *formicoides* |
| casent0903245 | Formicinae | *Myrmoteras* | *arcoelinae* |
| casent0903253 | Formicinae | *Myrmecorhynchus* | *carteri* |
| casent0903254 | Formicinae | *Myrmecorhynchus* | *nitidus* |
| casent0903259 | Formicinae | *Gigantiops* | *destructor* |
| casent0903286 | Formicinae | *Formica* | *podzolica* |
| casent0903597 | Formicinae | *Colobopsis* | *badia* |
| casent0903671 | Dorylinae | *Acanthostichus* | *lattkei* |
| casent0903772 | Dorylinae | *Cerapachys* | *sulcinodis* |
| casent0903830 | Ectatomminae | *Rhytidoponera* | *anceps* |
| casent0903924 | Ponerinae | *Myopias* | *bidens* |
| casent0903946 | Ponerinae | *Leptogenys* | *acutangula* |
| casent0904061 | Myrmicinae | *Manica* | *invidia* |
| casent0904544 | Myrmicinae | *Vollenhovia* | *brevicornis* |
| casent0904546 | Myrmicinae | *Vollenhovia* | *butteli* |
| casent0904573 | Myrmicinae | *Chelaner* | *aper* |
| casent0904585 | Myrmicinae | *Oxyepoecus* | *mandibularis* |
| casent0904614 | Myrmicinae | *Xenomyrmex* | *floridanus* |
| casent0904616 | Myrmicinae | *Tranopelta* | *subterranea* |
| casent0904647 | Myrmicinae | *Lophomyrmex* | *bedoti* |

| | | | |
|---|---|---|---|
| casent0904686 | Myrmicinae | *Meranoplus* | *armatus* |
| casent0904688 | Myrmicinae | *Myrmecina* | *americana* |
| casent0904863 | Myrmicinae | *Strongylognathus* | *afer* |
| casent0904984 | Myrmicinae | *Kalathomyrmex* | *emeryi* |
| casent0905008 | Myrmicinae | *Atta* | *cephalotes* |
| casent0905018 | Dolichoderinae | *Dolichoderus* | *affinis* |
| casent0905041 | Aneuretinae | *Aneuretus* | *simoni* |
| casent0905045 | Dolichoderinae | *Iridomyrmex* | *anceps* |
| casent0905654 | Formicinae | *Pseudolasius* | *binghami* |
| casent0905658 | Formicinae | *Pseudolasius* | *amaurops* |
| casent0905660 | Formicinae | *Euprenolepis* | *procera* |
| casent0905981 | Dorylinae | *Aenictus* | *aitkenii* |
| casent0906198 | Myrmicinae | *Nesomyrmex* | *pleuriticus* |
| casent0906264 | Formicinae | *Lepisiota* | *arabica* |
| casent0906283 | Formicinae | *Myrmoteras* | *barbouri* |
| casent0906285 | Formicinae | *Myrmoteras* | *bakeri* |
| casent0906288 | Formicinae | *Gesomyrmex* | *kalshoveni* |
| casent0906295 | Formicinae | *Cataglyphis* | *albicans* |
| casent0906313 | Formicinae | *Iberoformica* | *subrufa* |
| casent0906352 | Dorylinae | *Ooceraea* | *biroi* |
| casent0906441 | Formicinae | *Tapinolepis* | *longitarsis* |
| casent0906833 | Amblyoponinae | *Stigmatomma* | *besucheti* |
| casent0906856 | Formicinae | *Calomyrmex* | *albopilosus* |
| casent0906916 | Formicinae | *Paratrechina* | *antsingy* |
| casent0907042 | Dorylinae | *Zasphinctus* | *cedaris* |
| casent0907049 | Dorylinae | *Parasyscia* | *arnoldi* |
| casent0907055 | Dorylinae | *Eburopone* | *wroughtoni* |
| casent0907057 | Dorylinae | *Ooceraea* | *australis* |
| casent0907061 | Dorylinae | *Lioponera* | *binodis* |
| casent0907073 | Myrmeciinae | *Myrmecia* | *aberrans* |
| casent0907126 | Ponerinae | *Platythyrea* | *angusta* |
| casent0907133 | Ectatomminae | *Acanthoponera* | *goeldii* |
| casent0907206 | Proceratiinae | *Proceratium* | *algiricum* |
| casent0907208 | Proceratiinae | *Probolomyrmex* | *brevirostris* |
| casent0907214 | Ponerinae | *Odontoponera* | *transversa* |
| casent0907216 | Ponerinae | *Diacamma* | *assamense* |
| casent0907234 | Ponerinae | *Ophthalmopone* | *ilgii* |
| casent0907237 | Ponerinae | *Neoponera* | *apicalis* |
| casent0907253 | Ponerinae | *Pseudoneoponera* | *havilandi* |
| casent0907275 | Ponerinae | *Mesoponera* | *australis* |
| casent0907287 | Ponerinae | *Pseudoponera* | *stigma* |
| casent0907444 | Myrmicinae | *Metapone* | *greeni* |
| casent0907456 | Pseudomyrmecinae | *Tetraponera* | *allaborans* |
| casent0907582 | Dolichoderinae | *Tapinoma* | *antarcticum* |
| casent0907607 | Leptanillinae | *Leptanilla* | *escheri* |
| casent0907708 | Myrmicinae | *Aphaenogaster* | *phalangium* |
| casent0907743 | Myrmicinae | *Messor* | *aciculatus* |
| casent0907759 | Myrmicinae | *Oxyopomyrmex* | *krueperi* |
| casent0908323 | Myrmicinae | *Rhopalomastix* | *rothneyi* |
| casent0908325 | Myrmicinae | *Myrmicaria* | *birmana* |

| | | | |
|---|---|---|---|
| casent0908335 | Myrmicinae | *Cardiocondyla* | *batesii* |
| casent0908387 | Myrmicinae | *Crematogaster* | *abstinens* |
| casent0908643 | Myrmicinae | *Crematogaster* | *aberrans* |
| casent0908682 | Myrmicinae | *Chelaner* | *antarcticus* |
| casent0908722 | Myrmicinae | *Trichomyrmex* | *aberrans* |
| casent0908792 | Myrmicinae | *Bondroitia* | *lujae* |
| casent0908829 | Myrmicinae | *Solenopsis* | *abjectior* |
| casent0908844 | Myrmicinae | *Xenomyrmex* | *picquarti* |
| casent0908849 | Myrmicinae | *Solenopsis* | *bicolor* |
| casent0908923 | Myrmicinae | *Mayriella* | *overbecki* |
| casent0908933 | Myrmicinae | *Meranoplus* | *bellii* |
| casent0908990 | Myrmicinae | *Poecilomyrma* | *myrmecodiae* |
| casent0908996 | Myrmicinae | *Vombisidris* | *jacobsoni* |
| casent0909016 | Myrmicinae | *Temnothorax* | *aeolius* |
| casent0909210 | Myrmicinae | *Ochetomyrmex* | *semipolitus* |
| casent0909213 | Myrmicinae | *Wasmannia* | *auropunctata* |
| casent0909214 | Myrmicinae | *Wasmannia* | *iheringi* |
| casent0909242 | Myrmicinae | *Procryptocerus* | *batesi* |
| casent0909255 | Myrmicinae | *Procryptocerus* | *adlerzi* |
| casent0909350 | Myrmicinae | *Colobostruma* | *alinodis* |
| casent0909357 | Myrmicinae | *Mycocepurus* | *smithii* |
| casent0909360 | Myrmicinae | *Myrmicocrypta* | *foreli* |
| casent0909385 | Myrmicinae | *Mycetarotes* | *parallelus* |
| casent0909422 | Myrmicinae | *Acromyrmex* | *echinatior* |
| casent0909437 | Myrmicinae | *Amoimyrmex* | *bruchi* |
| casent0909446 | Myrmicinae | *Atta* | *bisphaerica* |
| casent0909495 | Dolichoderinae | *Liometopum* | *lindgreeni* |
| casent0909497 | Dolichoderinae | *Turneria* | *dahlii* |
| casent0909499 | Dolichoderinae | *Iridomyrmex* | *agilis* |
| casent0909525 | Dolichoderinae | *Ochetellus* | *epinotalis* |
| casent0909561 | Dolichoderinae | *Ochetellus* | *flavipes* |
| casent0909577 | Dolichoderinae | *Arnoldius* | *pusillus* |
| casent0909579 | Dolichoderinae | *Chronoxenus* | *dalyi* |
| casent0909582 | Dolichoderinae | *Azteca* | *aesopus* |
| casent0909712 | Dolichoderinae | *Forelius* | *albiventris* |
| casent0909720 | Dolichoderinae | *Forelius* | *breviscapus* |
| casent0909721 | Dolichoderinae | *Forelius* | *brasiliensis* |
| casent0909729 | Dolichoderinae | *Dorymyrmex* | *antarcticus* |
| casent0909743 | Dolichoderinae | *Dorymyrmex* | *antillanus* |
| casent0909806 | Dolichoderinae | *Tapinoma* | *albinase* |
| casent0909816 | Formicinae | *Melophorus* | *aeneovirens* |
| casent0909820 | Formicinae | *Prolasius* | *mjoebergellus* |
| casent0909828 | Formicinae | *Lasiophanes* | *picinus* |
| casent0909829 | Formicinae | *Lasiophanes* | *hoffmanni* |
| casent0909835 | Formicinae | *Notoncus* | *ectatommoides* |
| casent0909850 | Formicinae | *Plagiolepis* | *abyssinica* |
| casent0909870 | Formicinae | *Lepisiota* | *arnoldi* |
| casent0909933 | Formicinae | *Stigmacros* | *australis* |
| casent0910758 | Formicinae | *Calomyrmex* | *impavidus* |
| casent0910970 | Formicinae | *Pseudolasius* | *amblyops* |

| | | | |
|---|---|---|---|
| casent0911002 | Formicinae | *Paraparatrechina* | *butteli* |
| casent0911016 | Formicinae | *Paraparatrechina* | *caledonica* |
| casent0911065 | Formicinae | *Bajcaridris* | *kraussii* |
| casent0911107 | Formicinae | *Cataglyphis* | *aenescens* |
| casent0911143 | Dorylinae | *Ooceraea* | *besucheti* |
| casent0911145 | Myrmicinae | *Cyphoidris* | *werneri* |
| casent0911369 | Dorylinae | *Cheliomyrmex* | *megalonyx* |
| casent0911453 | Leptanillinae | *Leptanilla* | *tanakai* |
| casent0911480 | Dolichoderinae | *Bothriomyrmex* | *corsicus* |
| casent0911493 | Dolichoderinae | *Chronoxenus* | *wroughtonii* |
| casent0912167 | Formicinae | *Alloformica* | *flavicornis* |
| casent0912169 | Formicinae | *Proformica* | *caucasea* |
| casent0912359 | Formicinae | *Tapinolepis* | *candida* |
| casent0912434 | Formicinae | *Stigmacros* | *barretti* |
| casent0912491 | Myrmicinae | *Atta* | *colombica* |
| casent0912506 | Myrmicinae | *Paramycetophylax* | *bruchi* |
| casent0912515 | Myrmicinae | *Sericomyrmex* | *amabilis* |
| casent0912524 | Myrmicinae | *Mycetomoellerius* | *haytianus* |
| casent0912540 | Myrmicinae | *Wasmannia* | *affinis* |
| casent0912875 | Myrmicinae | *Atopomyrmex* | *mocquerysi* |
| casent0913090 | Myrmicinae | *Pogonomyrmex* | *abdominalis* |
| casent0913152 | Myrmicinae | *Messor* | *alexandri* |
| casent0913240 | Myrmicinae | *Oxyopomyrmex* | *emeryi* |
| casent0913243 | Myrmicinae | *Oxyopomyrmex* | *insularis* |
| casent0913491 | Myrmicinae | *Carebara* | *alluaudi* |
| casent0913643 | Formicinae | *Proformica* | *coriacea* |
| casent0913651 | Myrmicinae | *Goniomma* | *blanci* |
| casent0913718 | Pseudomyrmecinae | *Tetraponera* | *aethiops* |
| casent0913754 | Formicinae | *Oecophylla* | *longinoda* |
| casent0913958 | Myrmicinae | *Dicroaspis* | *laevidens* |
| casent0913959 | Myrmicinae | *Lordomyrma* | *azumai* |
| casent0913969 | Myrmicinae | *Stenamma* | *africanum* |
| casent0914048 | Myrmicinae | *Myrmicaria* | *baumi* |
| casent0914059 | Myrmicinae | *Megalomyrmex* | *balzani* |
| casent0914065 | Myrmicinae | *Dilobocondyla* | *borneensis* |
| casent0914317 | Myrmicinae | *Monomorium* | *abeillei* |
| casent0914340 | Ponerinae | *Ophthalmopone* | *hottentota* |
| casent0914459 | Myrmicinae | *Pogonomyrmex* | *occidentalis* |
| casent0914500 | Ponerinae | *Simopelta* | *andersoni* |
| casent0914503 | Ponerinae | *Simopelta* | *laevigata* |
| casent0914576 | Myrmicinae | *Crematogaster* | *nigropilosa* |
| casent0914580 | Formicinae | *Rossomyrmex* | *anatolicus* |
| casent0914667 | Myrmicinae | *Epopostruma* | *areosylva* |
| casent0914886 | Myrmicinae | *Basiceros* | *conjugans* |
| casent0914888 | Myrmicinae | *Basiceros* | *manni* |
| casent0914891 | Myrmicinae | *Eurhopalothrix* | *alopeciosa* |
| casent0914907 | Myrmicinae | *Octostruma* | *amrishi* |
| casent0914955 | Myrmicinae | *Atopomyrmex* | *cryptoceroides* |
| casent0914984 | Myrmicinae | *Nesomyrmex* | *anduzei* |
| casent0915106 | Ectatomminae | *Ectatomma* | *edentatum* |

| | | | |
|---|---|---|---|
| casent0915183 | Ponerinae | *Emeryopone* | *franzi* |
| casent0915184 | Ponerinae | *Emeryopone* | *loebli* |
| casent0915247 | Ponerinae | *Ectomomyrmex* | *aequalis* |
| casent0915248 | Ponerinae | *Brachyponera* | *arcuata* |
| casent0915269 | Ponerinae | *Loboponera* | *nasica* |
| casent0915286 | Ponerinae | *Plectroctena* | *dentata* |
| casent0915294 | Ponerinae | *Ponera* | *coarctata* |
| casent0915323 | Pseudomyrmecinae | *Pseudomyrmex* | *adustus* |
| casent0915330 | Dorylinae | *Cylindromyrmex* | *longiceps* |
| casent0915337 | Dolichoderinae | *Dorymyrmex* | *baeri* |
| casent0915348 | Ectatomminae | *Ectatomma* | *gibbum* |
| casent0915350 | Ectatomminae | *Typhlomyrmex* | *meire* |
| casent0915407 | Myrmeciinae | *Myrmecia* | *fulvipes* |
| casent0915448 | Myrmicinae | *Goniomma* | *collingwoodi* |
| casent0915488 | Ponerinae | *Hypoponera* | *abeillei* |
| casent0915627 | Formicinae | *Polyergus* | *bicolor* |
| casent0915629 | Formicinae | *Alloformica* | *aberrans* |
| casent0915657 | Ponerinae | *Mayaponera* | *constricta* |
| casent0915664 | Ponerinae | *Hagensia* | *havilandi* |
| casent0915697 | Myrmicinae | *Basiceros* | *convexiceps* |
| casent0915705 | Formicinae | *Anoplolepis* | *fallax* |
| casent0915715 | Formicinae | *Lepisiota* | *capensis* |
| casent0915930 | Ectatomminae | *Heteroponera* | *carinifrons* |
| casent0915936 | Ponerinae | *Belonopelta* | *attenuata* |
| casent0916034 | Myrmicinae | *Allomerus* | *octoarticulatus* |
| casent0916035 | Myrmicinae | *Diplomorium* | *longipenne* |
| casent0916097 | Ponerinae | *Diacamma* | *aureovestitum* |
| casent0916796 | Amblyoponinae | *Stigmatomma* | *amblyops* |
| casent0917031 | Myrmicinae | *Pristomyrmex* | *africanus* |
| casent0917203 | Formicinae | *Colobopsis* | *aruensis* |
| casent0917214 | Myrmicinae | *Harpagoxenus* | *zaisanicus* |
| casent0917363 | Myrmicinae | *Perissomyrmex* | *monticola* |
| casent0917433 | Myrmicinae | *Strongylognathus* | *arnoldii* |
| casent0917680 | Myrmicinae | *Myrmica* | *ademonia* |
| casent0917752 | Myrmicinae | *Messor* | *aegyptiacus* |
| casent0919619 | Myrmicinae | *Aretidris* | *buenaventei* |
| casent0919731 | Myrmicinae | *Cardiocondyla* | *brachyceps* |
| casent0919746 | Myrmicinae | *Leptothorax* | *athabasca* |
| casent0919792 | Myrmicinae | *Novomessor* | *albisetosus* |
| casent0919967 | Myrmicinae | *Trachymyrmex* | *arizonensis* |
| casent0919970 | Myrmicinae | *Paratrachymyrmex* | *intermedius* |
| casent0919976 | Myrmicinae | *Trachymyrmex* | *carinatus* |
| casent0922019 | Myrmicinae | *Acromyrmex* | *ambiguus* |
| casent0922036 | Myrmicinae | *Apterostigma* | *bolivianum* |
| casent0922150 | Myrmicinae | *Xerolitor* | *explicatus* |
| casent0922224 | Myrmicinae | *Orectognathus* | *chyzeri* |
| casent0922225 | Myrmicinae | *Orectognathus* | *biroi* |
| casent0922434 | Ponerinae | *Ectomomyrmex* | *aciculatus* |
| casent0922626 | Myrmicinae | *Procryptocerus* | *balzani* |
| casent0922687 | Myrmicinae | *Aphaenogaster* | *aktaci* |

| | | | |
|---|---|---|---|
| casent0922741 | Myrmicinae | *Manica* | *hunteri* |
| casent0922778 | Myrmicinae | *Myrmica* | *aimonissabaudiae* |
| casent0922897 | Myrmicinae | *Vombisidris* | *acherdos* |
| casent0922901 | Myrmicinae | *Lenomyrmex* | *mandibularis* |
| casent0922947 | Ponerinae | *Leptogenys* | *academica* |
| casent0922971 | Myrmicinae | *Myrmecina* | *australis* |
| casent0922982 | Myrmicinae | *Pristomyrmex* | *bispinosus* |
| casent0923015 | Myrmicinae | *Hylomyrma* | *dentiloba* |
| casent0923154 | Ponerinae | *Bothroponera* | *cariosa* |
| castype00622 | Myrmicinae | *Novomessor* | *cockerelli* |
| castype05026 | Myrmicinae | *Stereomyrmex* | *dispar* |
| castype06888 | Ectatomminae | *Acanthoponera* | *minor* |
| castype09451 | Ponerinae | *Simopelta* | *laticeps* |
| castype11766 | Dolichoderinae | *Chronoxenus* | *rossi* |
| ecofog-it14-0797-30 | Myrmicinae | *Mycetomoellerius* | *farinosus* |
| ecofog-kw14-0023-12 | Myrmicinae | *Oxyepoecus* | *ephippiatus* |
| ecofog-tr17-0175-08 | Myrmicinae | *Mycetarotes* | *acutus* |
| fmnhins0000051220 | Ponerinae | *Cryptopone* | *butteli* |
| focol0340-1 | Dorylinae | *Lioponera* | *bicolor* |
| focol0376 | Dorylinae | *Chrysapace* | *sauteri* |
| focol0535 | Dolichoderinae | *Azteca* | *adrepens* |
| focol0554 | Formicinae | *Rossomyrmex* | *minuchae* |
| focol0952 | Ponerinae | *Pachycondyla* | *impressa* |
| focol0967 | Ponerinae | *Austroponera* | *castanea* |
| focol1073 | Ponerinae | *Odontomachus* | *aciculatus* |
| focol1569 | Myrmicinae | *Ancyridris* | *rupicapra* |
| focol1945 | Myrmicinae | *Myrmecina* | *bandarensis* |
| focol1970 | Myrmicinae | *Dilobocondyla* | *cataulacoidea* |
| focol2001 | Myrmicinae | *Podomyrma* | *abdominalis* |
| focol2183 | Myrmicinae | *Ocymyrmex* | *barbiger* |
| focol2219 | Formicinae | *Notoncus* | *gilberti* |
| focol2561 | Formicinae | *Echinopla* | *arfaki* |
| focol2562 | Formicinae | *Echinopla* | *densistriata* |
| focol2809 | Dolichoderinae | *Anonychomyrma* | *angusta* |
| inb0003214454 | Myrmicinae | *Lenomyrmex* | *colwelli* |
| inb0003660648 | Ponerinae | *Rasopone* | *cryptergates* |
| inb0003679758 | Myrmicinae | *Lachnomyrmex* | *laticeps* |
| inbiocri001279879 | Myrmicinae | *Adelomyrmex* | *longinoi* |
| jdm32-002000-1 | Formicinae | *Melophorus* | *attenuipes* |
| jtlc000002756 | Formicinae | *Myrmelachista* | *joycei* |
| jtlc000003150 | Myrmicinae | *Adelomyrmex* | *betoi* |
| jtlc000005275 | Formicinae | *Myrmelachista* | *amicta* |
| jtlc000005879 | Myrmicinae | *Stenamma* | *alas* |
| jtlc000015205 | Ponerinae | *Pseudoponera* | *gilberti* |
| mb619-2 | Myrmicinae | *Strongylognathus* | *caeciliae* |
| mcz121421139 | Myrmicinae | *Sericomyrmex* | *lutzi* |
| ncbs-av849 | Myrmicinae | *Tyrannomyrmex* | *alii* |
| okent0035688 | Leptanillinae | *Protanilla* | *lini* |
| psw11581-5 | Myrmicinae | *Perissomyrmex* | *snyderi* |

| | | | |
|---|---|---|---|
| psw7668-22 | Myrmicinae | *Xenomyrmex* | *panamanus* |
| qmt120169 | Myrmicinae | *Adlerzia* | *froggatti* |
| qmt162820 | Formicinae | *Teratomyrmex* | *greavesi* |
| rmcaent000017740 | Ponerinae | *Anochetus* | *africanus* |
| sam-hym-c002831 | Ponerinae | *Psalidomyrmex* | *procerus* |
| sam-hym-c008788b | Ponerinae | *Bothroponera* | *cavernosa* |
| ufv-labecol-000105 | Ponerinae | *Thaumatomyrmex* | *contumax* |
| ufv-labecol-001083 | Myrmicinae | *Acromyrmex* | *ameliae* |
| ufv-labecol-004945 | Myrmicinae | *Mycetophylax* | *auritus* |
| ufv-labecol-009622 | Proceratiinae | *Discothyrea* | *horni* |
| usnment00445715 | Myrmicinae | *Acanthognathus* | *lentus* |
| usnment00529071 | Myrmicinae | *Veromessor* | *andrei* |
| usnment00755089 | Formicinae | *Nylanderia* | *acuminata* |
| usnment00755129 | Formicinae | *Prenolepis* | *darlena* |
| usnment00757177 | Formicinae | *Brachymyrmex* | *attenuatus* |
| usnment00758173 | Myrmicinae | *Cyatta* | *abscondita* |
| usnment00921167 | Myrmicinae | *Mycetophylax* | *asper* |
| usnment01125207 | Myrmicinae | *Sericomyrmex* | *bondari* |
| usnment01126300 | Dorylinae | *Syscia* | *honduriana* |
| antweb1041405 | Ponerinae | *Pachycondyla* | *crassinoda* |
| casent0006154 | Myrmicinae | *Lordomyrma* | *crawleyi* |
| casent0102245 | Myrmicinae | *Terataner* | *balrog* |
| casent0249450 | Dorylinae | *Cheliomyrmex* | *andicola* |
| casent0250350 | Ponerinae | *Platythyrea* | *arnoldi* |
| casent0254323 | Ponerinae | *Boloponera* | *ikemkha* |
| casent0814561 | Ponerinae | *Parvaponera* | *darwinii* |
| casent0911330 | Dorylinae | *Dorylus* | *aggressor* |
| casent0912513 | Myrmicinae | *Myrmicocrypta* | *urichi* |
| casent0914439 | Formicinae | *Camponotus* | *acutirostris* |
